# Activation of the impaired NAMPT/SIRT7/SOD2 axis restores alveolar progenitor cell homeostasis in idiopathic pulmonary fibrosis and reverses pulmonary fibrosis in mice

**DOI:** 10.1101/2025.06.30.662456

**Authors:** Xuexi Zhang, Xue Liu, Yujie Qiao, Anas Rabata, Ningshan Liu, Changfu Yao, Tanyalak Parimon, Danica Chen, Peter Chen, Barry Stripp, Stephen J. Gardell, Dianhua Jiang, Paul W. Noble, Jiurong Liang

## Abstract

Alveolar type II (AT2) progenitor cell exhaustion and impaired regenerative capacity are key pathogenic hallmarks in idiopathic pulmonary fibrosis (IPF). Nicotinamide adenine dinucleotide (NAD^+^) functions as a central regulator of cellular energy metabolism. We have reported that downregulation of NAD^+^-dependent sirtuin signaling contributes to the impaired progenitor function of IPF AT2s. In this study, we identified that a key NAD^+^ biosynthesis enzyme, nicotinamide phosphoribosyltransferase (NAMPT), is significantly downregulated in IPF AT2s. NAMPT deficiency impairs AT2 renewal and enhances lung fibrosis through downregulation of SIRT7 and SOD2, which results in increased oxidative stress, mitochondrial dysfunction, induction of pathological transitional gene expression and impaired regenerative capacity to generate alveolar type I (AT1) cell required for gas exchange. Mice with deletion of Nampt in AT2s showed severely impaired AT2 renewal and increased susceptibility to bleomycin lung injury and spontaneous fibrois. Activation of NAMPT by small molecule activators promoted AT2 renewal, restored homeostasis, and reversed lung fibrosis. NAMPT activation could be a therapeutic strategy for restoring AT2 progenitor function and halting or reversing progressive pulmonary fibrosis.

## Introduction

Idiopathic pulmonary fibrosis (IPF) is a fatal form of lung interstitial lung disease with limited treatment options^1–3^. The median survival of IPF patients from diagnosis is 2–4 years^3^. Despite two FDA-approved treatments with limited therapeutic effect in restoring lung functions^4,5^, IPF remains the leading cause of lung transplant. It is believed that loss of epithelial integrity due to repeated epithelial cell injury and inadequate alveolar epithelial repair is a seminal causal event in the pathogenesis of IPF^2,6–12^. Type 2 alveolar epithelial cells (AT2s) function as progenitor cells to maintain alveolar epithelial integrity during homeostasis and in response to injury^13–15^. The AT2 cell population in IPF lungs is decreased and the remaining AT2s in the diseased lungs have impaired renewal capacity and fail to generate type I alveolar epithelial cells (AT1s) that are essential for gas exchange^6,8,16^. However, the molecular mechanisms that control AT2 progenitor cell renewal and unremitting lung fibrosis remain poorly understood.

Nicotinamide adenine dinucleotides (NAD^+^ and NADH) are central regulators of cell energy metabolism^17,18^. Published studies showed that NAD^+^ levels are decreased in IPF lungs^19,20^, and NAD^+^ deficiency is associated with lung fibrosis^19–21^. Furthermore, NAD^+^ levels have been shown to decrease with aging^22,23^. We have recently reported that NAD^+^ dependent sirtuin signaling is crucial for AT2 progenitor cell renewal. However, the causes for the reduced NAD^+^ level in AT2s from fibrotic lungs has not been elucidated.

NAD^+^ biosynthesis is synthesized via the *de novo,* Preiss-Handler, and salvage pathways which use tryptophan, nicotinic acid and nicotinamide (NAM), respectively, as the precursor for the pyridine ring structure^22,24,25^. The salvage pathway is the major contributor to NAD^+^ synthesis in a wide variety of tissues^24,26^. The putative rate-determining step for the salvage pathway is nicotinamide phosphoribosyltransferase (NAMPT) which produces nicotinamide mononucleotide (NMN). NAD^+^ is formed from NMN by NMN adenylytransferase (NMNAT)^22,24^. NAMPT has been implicated in regulating stem cell functions, mitochondrial bioenergetics, and cell apoptosis and senescence in neurodegenerative diseases, endothelial cells and human iPS cells^27–31^. NAMPT-mediated NAD^+^ biosynthesis protects against CCl4-induced liver fibrosis^32^. However, the role of NAD^+^ metabolism and NAMPT activity in regulating AT2 progenitor cell function in in IPF and experimental lung fibrosis has not been fully investigated.

NAD^+^ is a cosubstrate for the sirtuin family (SIRT1-7)^22,24^, a family of NAD^+^-dependent deacylases which have been implicated in promoting stem cell function and cell rejuvenation^18,33,34^. We reported that downregulation of NAD^+^-dependent sirtuin signaling contributed to the impaired progenitor function of IPF AT2s, and NAD^+^ precursors promoted AT2 renewal^8^. Studies showed that SIRT7 plays an important role in regulating cellular energy consumption^35^ and maintaining mitochondrial homeostasis^36,37^. SIRT7 expression was reduced in aging^38–40^, whereas upregulation of SIRT7 improved regeneration of aged hematopoietic stem cells^38^. SIRT7 protected against cellular senescence in mammalian cell lines^41^ and in mesenchymal stem cells^42^. Hepatic SIRT7 suppressed endoplasmic reticulum (ER) stress and prevented fatty liver disease^43^. The NAD^+^ precursor, nicotinamide mononucleotide (NMN), promoted intestinal stem cells, reduced cellular reactive oxygen species (ROS) levels, and protected against radiation injury by enhancing SIRT6 and SIRT7 activities^44^. However, the role of NAD^+^ and NAMPT associated SIRT7 activity in regulating AT2 progenitor activity is unknown.

NAD^+^ is a ubiquitous enzyme cofactor that is obligatory for cellular redox reactions. Reduced levels of NAD^+^ are linked to dysregulation of the oxidative stress responses and mitochondrial dysfunction in aging and aging-associated diseases^45,46^. Epithelial cells in IPF lungs showed increased oxidative stress^47–51^ and diminished expression of oxidative response proteins^52–54^, leading to senescence and apoptosis. SOD2 protects alveolar epithelial cells from lung injury^55,56^. SIRT7 was shown to promote cell proliferation, inhibit apoptosis and oxidative stress, and maintain mitochondrial homeostasis^57–59^. The interaction between SIRT7 and SOD2 in AT2 progenitor cells has not been investigated.

The pivotal role of NAD^+^ and the therapeutic promise of preserving tissue NAD^+^ levels have created keen interest in the NAD^+^ precursors, NMN, nicotinamide (NAM) and nicotinamide riboside (NR)^25^. A recent addition to the NAD^+^ booster arsenal are small molecules (NAMPT activators) that bind directly to NAMPT and promote NMN synthesis^28,60^.

In this study, we discovered that the key NAD^+^ biosynthesis enzyme, NAMPT, was significantly down regulated in IPF AT2s. We further determined that both genetic and pharmacologic activation of NAMPT promoted, whereas NAMPT inhibition suppressed, AT2 progenitor renewal in 3D organoid cultures and in mice in vivo. Our pursuit of the molecular mechanisms by which NAMPT regulates AT2 progenitor cell renewal revealed that NAMPT promotes AT2 progenitor cell renewal through SIRT7 and SOD2. We hypothesize that NAMPT deficiency in IPF AT2s downregulates SIRT7 and SOD2 making these cells more vulnerable to oxidative stress and mitochondrial dysfunction, ultimately eliciting impaired AT2 cell renewal and severe lung fibrosis. Interestingly, treatment with NAMPT activator increased AT2 recovery and attenuated bleomycin induced lung fibrosis in vivo with both young and old mice. To our knowledge, this is the first evidence showing that the NAMPT/SIRT7/SOD2 axis plays a critical role in regulating AT2 renewal and lung fibrosis, and suggests that we have identified the “Achilles Heel” of IPF.

## Results

### Decreased expression of a NAD^+^ biosynthesis enzyme, NAMPT, in IPF AT2s

Reports have shown that NAD^+^ is decreased in IPF lung tissues^19,20^. The salvage pathway involving NAMPT is the major route for NAD^+^ biosynthesis in the lung^26^. scRNA-seq analysis using flow enriched human lung epithelial cells from both healthy and IPF lungs^8^ revealed that NAMPT is highly expressed in AT2s (Figure 1A and S1A). Interestingly, the expression levels of NAMPT were dramatically lower in IPF AT2s (Figure 1B and 1C). Decreased NAMPT expression in IPF AT2s was also evident in multiple recently published scRNA-seq data sets^61–64^ (Figure 1D). Using freshly isolated AT2s from healthy and IPF lungs, we further showed decreased NAMPT mRNA levels in IPF AT2s with qPCR (Figure 1E) as well as decreased NAMPT protein levels via both single cell western blot (Figure 1F and S1B) and conventional western blot (Figure 1G). With immunofluorescence staining, we demonstrated that NAMPT was co-localized with the human AT2 marker HTII-280 (Figure 1H) and the fluorescence intensity of NAMPT staining in AT2s of IPF lung sections was decreased compared to that in the AT2s of healthy lung sections (Figure 1H and 1I).

**Figure 1.**
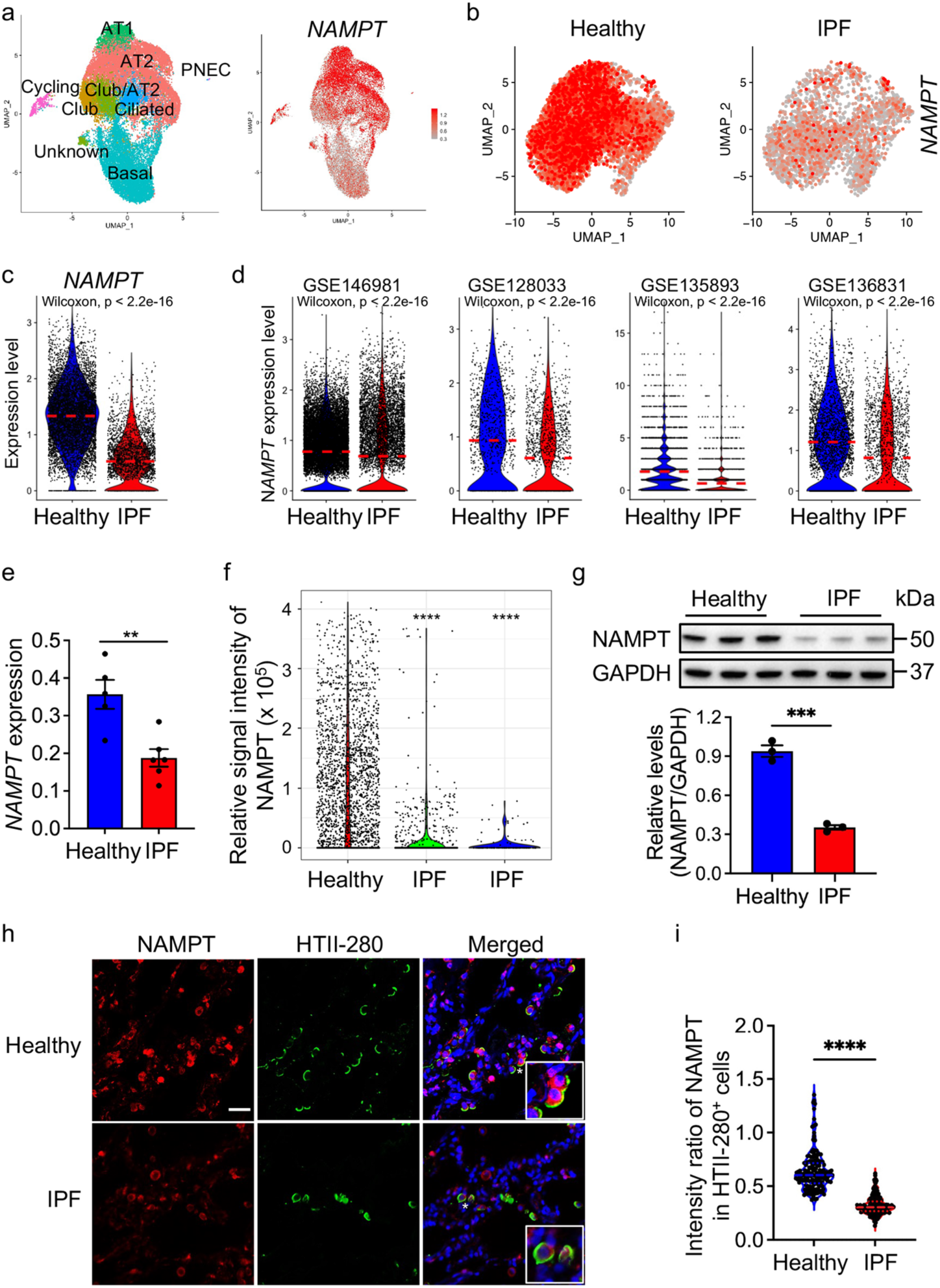
Decreased expression of a NAD^+^ biosynthesis enzyme, NAMPT, in IPF AT2s. (A) UMAP visualization of epithelial cell types and *NAMPT* expression in healthy and IPF lung from in house scRNA-seq dataset. (B) UMAP visualization of *NAMPT* expression in AT2s from healthy and IPF lungs. (C and D) Violin plots showing *NAMPT* expression levels in AT2s from in house scRNA-seq and recently published scRNA-seq datasets. (E) Real-time PCR analysis of *NAMPT* expression in freshly isolated AT2s from healthy and IPF lungs (N = 5-6 per group, **p < 0.01 by unpaired t test). (F) Single-cell western blot analysis of NAMPT expression in freshly isolated AT2s from healthy and IPF lungs, with β-Tubulin as a loading control (****p < 0.0001 by ANOVA). (G) Western blot analysis and quantification of NAMPT expression in immortalized AT2s from healthy and IPF lungs. GAPDH served as loading control. ***p < 0.001 by unpaired t test. (H) Co-staining of NAMPT (red) and HTII-280 (green, an AT2 cell marker) on healthy and IPF lung sections. The staining was performed with lungs sections from three IPF patients and three healthy donors. Representative cells (asterisks) are shown at higher magnification. Scale bars, 20 μm. (I) Quantification of NAMPT staining (relative signal intensity) in individual AT2s (HTII-280^+^), shown as violin plots with median and quartiles (50-60 cells/section, 3 subjects/group, ****p < 0.0001 by unpaired t test). Data are shown as the mean ± SEM. See also Figure S1

We also compared NAMPT expression in fibroblasts from fibrotic lung tissues with integrated scRNA-seq data analysis. The expression levels of NAMPT were not significantly different between human fibroblasts from healthy and IPF lungs (Figure S1C), and between mouse fibroblasts from bleomycin induced fibrotic lungs and control lungs (Figure S1D).

### NAMPT regulates AT2 progenitor cell regeneration

Next, we tested the role of NAMPT in regulating AT2 progenitor cell regeneration. Small molecule NAMPT activators (NAT, NAT-5r and SBI-797812) were added to 3D organoid cultures of primary AT2s freshly isolated from healthy and IPF lungs and colony forming efficiency (CFE) was determined. NAT^28^ increased CFE of AT2s from both healthy and IPF lungs in a dose dependent manner (Figure 2A and 2B). To test if NAT was exerting its effect via NAMPT, we knocked out NAMPT (NAMPT^KO^) in immortalized human AT2s with CRISPR/Cas9 (Figure S2A). NAT treatment increased the NAD^+^/NADH ratio in control cells but not in the cells with NAMPT knockout (Figure S2B), pointing to NAMPT being the target of NAT action. Notably, two other NAMPT activators, SBI-797812^60^ and NAT-5r^28^ also promoted AT2 renewal (Figure 2C and 2D). Since NAT exhibited high NAMPT activation at low concentrations of NAD^+^ precursor^28^, we used NAT in the further studies.

**Figure 2.**
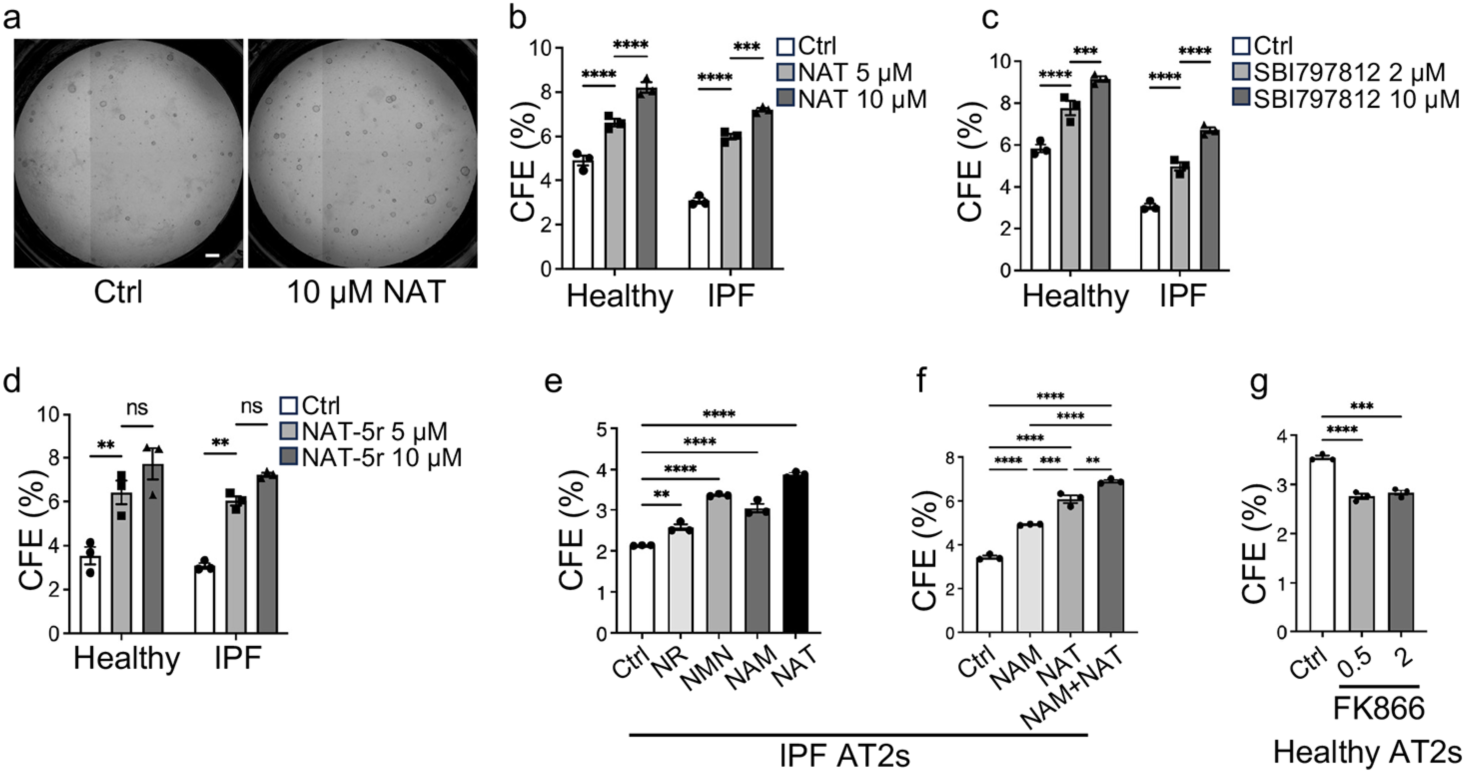
NAMPT regulates AT2 progenitor cell regeneration. (A) Representative images of human AT2 colonies cultured with NAT or DMSO control. Scale bars, 300 μm. (B-D) Colony forming efficiency (CFE) in 3D organoid cultures of AT2s from healthy and IPF lungs treated with the NAMPT activators (B) NAT, or (C) SBI797812, and (D) NAT-5r (n = 3, ns, no significant, **p < 0.01, ***p < 0.001, ****p < 0.0001 by ANOVA). (E and F) Colony forming efficiency (CFE) in 3D organoid cultures of AT2s from IPF lungs treated with NAMPT precursors NR, NMN and NAM, as well as NAMPT activators NAT (n = 3 per group, **p < 0.01, ***p < 0.001, ****p < 0.0001 by ANOVA). (G) CFE in 3D organoid cultures of AT2s from healthy lungs treated with the NAMPT inhibitor FK866 at different concentration (in nM) or vehicle control (n = 3, ****p < 0.0001 by ANOVA.). Data are shown as the mean ± SEM. See also Figure S2

To determine if NAD^+^ precursors could replicate the effects of NAT, we compared the effects of NR, NMN, and NAM, with NAT in 3D organoid cultures. Our results indicate that NAT has the strongest effect on IPF AT2 cell renewal (Figure 2E). Given that the NAMPT substrate NAM may also be reduced in IPF AT2s, we tested NAM alone and in combination with NAT in 3D organoid cultures. The combination of NAT and NAM exhibited a greater effect on IPF AT2 regenerative capacity (Figure 2F). In contrast, blocking NAMPT activity with the highly potent and selective NAMPT inhibitor, FK866, suppressed renewal of human AT2s (Figure 2G), which further confirmed the role of NAMPT in regulating AT2 progenitor function. Additionally, neither NAT nor FK866 had a significant effect on AT2 cell growth (Fig. S2C and S2D).

### NAMPT regulates SOD2 expression and oxidative stress responses of AT2s

To investigate the molecular mechanisms by which NAMPT regulates AT2 progenitor function, we separated human AT2s from healthy and IPF lungs by their *NAMPT* expression levels into NAMPT high (NAMPT^hi^, expression levels >1) and NAMPT low (NAMPT^low^, expression levels <=1) AT2s (Figure 3A) and identified differentially expressed genes (DEGs). The most downregulated genes in NAMPT^low^ AT2s was superoxide dismutase 2 (*SOD2*) (Figure 3B and 3C). Other oxidative stress response genes including *HIF1A* and *TXNRD1* along with AT2 marker genes *SLC34A2* and *SFTPC* were also downregulated in NAMPT^low^ AT2s (Figure 3B). Importantly, SOD2 and multiple oxidative stress response genes^65–68^ including *NQO1*, *TXNRD1*, and *ROMO1* were downregulated in IPF AT2s compared to AT2s from healthy donor lungs (Figure 3D). Consistent with these transcriptomic results, we further showed that IPF AT2s displayed increased mitochondrial superoxide levels compared to healthy AT2s as assayed by flow cytometry (Figure 3E and 3F).

**Figure 3.**
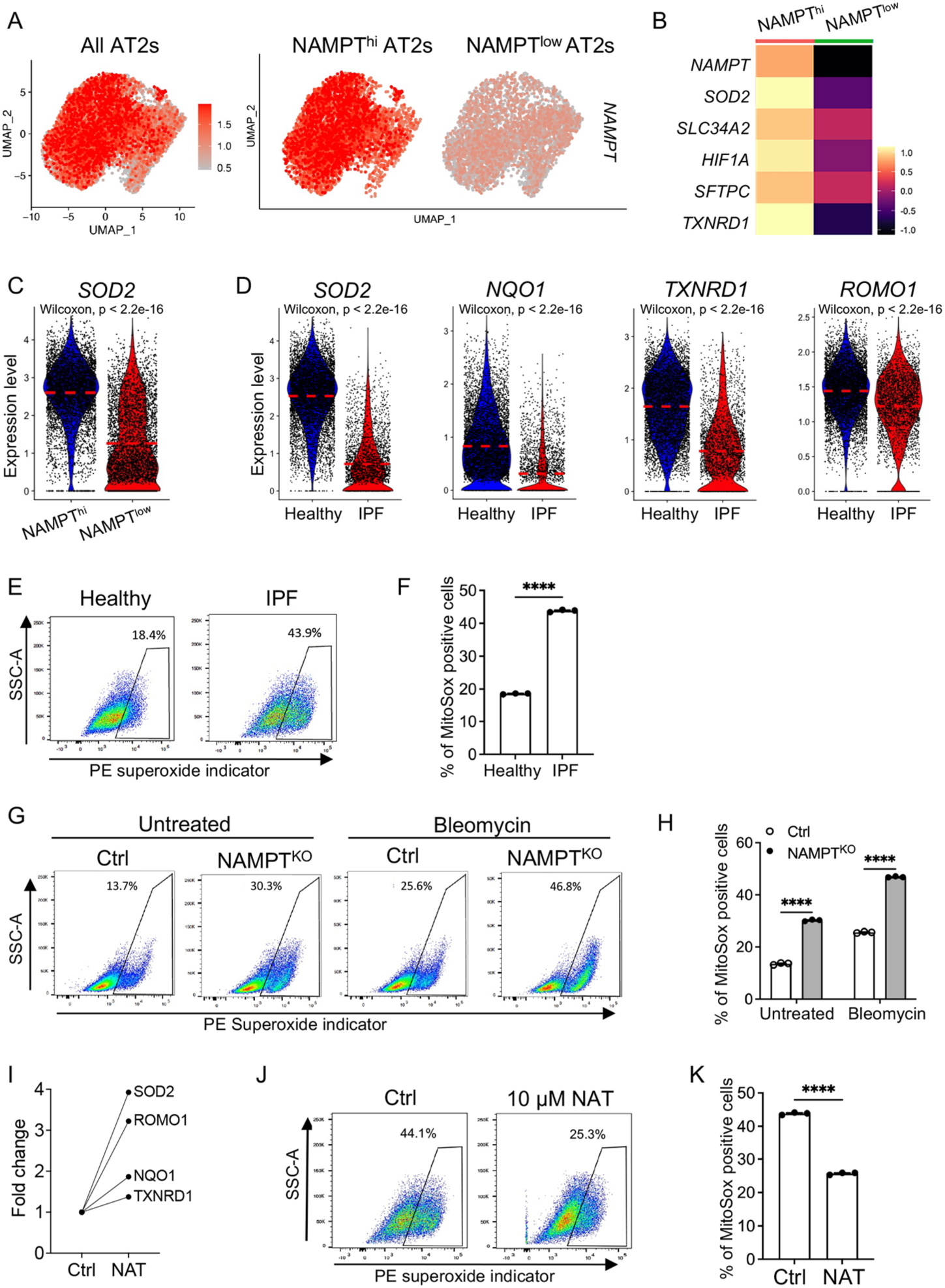
NAMPT regulates SOD2 expression and oxidative stress response of AT2s. (A) UMAP visualization of *NAMPT* expression in total, NAMPT^hi^ and NAMPT^low^ human AT2s. (B) Heatmap of differentially expressed genes (DEGs) in NAMPT^hi^ versus NAMPT^low^ human AT2s. (C) Violin plots showing *SOD2* expression levels in NAMPT^hi^ and NAMPT^low^ human AT2s. (D) Violin plots showing the expression levels of *SOD2*, *NQO1*, *TXNRD1*, and *ROMO1* in healthy and IPF AT2s from in house scRNA-seq dataset. (E and F) Flow cytometry analysis of mitochondrial superoxide levels in immortalized AT2s from healthy and IPF lungs. (E) Gating strategy and (F) percent of mitochondrial superoxide high cells (n = 3 per group, ****p < 0.0001 by unpaired t test). (G and H) Flow cytometry analysis of mitochondrial superoxide levels in NAMPT^KO^ and control immortalized AT2s, with and without 100 μg/ml bleomycin treated treatment. (G) Gating strategy and (H) percent of mitochondrial superoxide high cells (n = 3 per group, ****p < 0.0001 by ANOVA). (I) Fold change in the expression of antioxidant genes in AT2s from IPF patients with NAT or DMSO, analyzed by bulk RNA-seq. (J and K) Flow cytometry analysis of mitochondrial superoxide levels in immortalized IPF AT2s treated with NAT or DMSO control. (J) Gating strategy and (K) percent of mitochondrial superoxide high cells (n = 3 per group, ****p < 0.0001 by unpaired t test). Data are shown as the mean ± SEM.

Next, we looked at the oxidative stress response of human AT2s with NAMPT knockout (Figure S2A). Knockout of NAMPT increased mitochondrial superoxide levels in AT2s both at baseline and after bleomycin treatment (Figure 3G and 3H). Importantly, activation of NAMPT by the activator NAT markedly increased expression of *SOD2* and other oxidative stress response genes in IPF AT2s (Figure 3I). NAT treatment also decreased mitochondrial superoxide levels in IPF AT2s (Figure 3J and 3K). These data suggest that NAMPT regulates oxidative responses of AT2s and NAMPT deficiency is associated with increased oxidative stress of IPF AT2s.

### NAMPT regulates SIRT7 expression and SIRT7 deacetylates SOD2 in AT2s

The enzyme activity of SOD2 was shown to be increased by deacetylation of lysine residues 68 (K68) and 122 (K122) situated at the SOD active site^69,70^. Sirtuins deacetylate SOD2 in an NAD^+^-dependent manner^70–72^. Using bulk RNA-seq analysis, we found that NAT treatment of isolated IPF AT2s dramatically increased *SIRT7* expression (Figure 4A). Other sirtuin family genes (*SIRT1*, *2*, *3*, *5*) were also increased but more modestly (Figure 4A). SIRT7 was highly expressed in healthy human AT2s, and its expression levels were significantly downregulated in AT2s from IPF lungs, both in terms of median expression levels (Figure 4B) and the percentages of SIRT7-positive cells (Figure 4C). We confirmed decreased SIRT7 protein levels in IPF AT2s with western blot analysis (Figure 4D). NAMPT activation (NAMPT^ACT^) in IPF AT2s by CRISPR/Cas9 increased SIRT7 protein levels (Figure 4E) while NAMPT knockout in healthy AT2s decreased SIRT7 expression and increased K122-acetylation-SOD2 (the inactive form of SOD2) (Figure 4F). Furthermore, knockout of SIRT7 (SIRT7^KO^) in immortalized human AT2s increased K122-acetylation-SOD2 levels, while total SOD2 expression was not affected (Figure 4G). These data suggested that SIRT7 deacetylates and activates SOD2 in AT2s.

**Figure 4.**
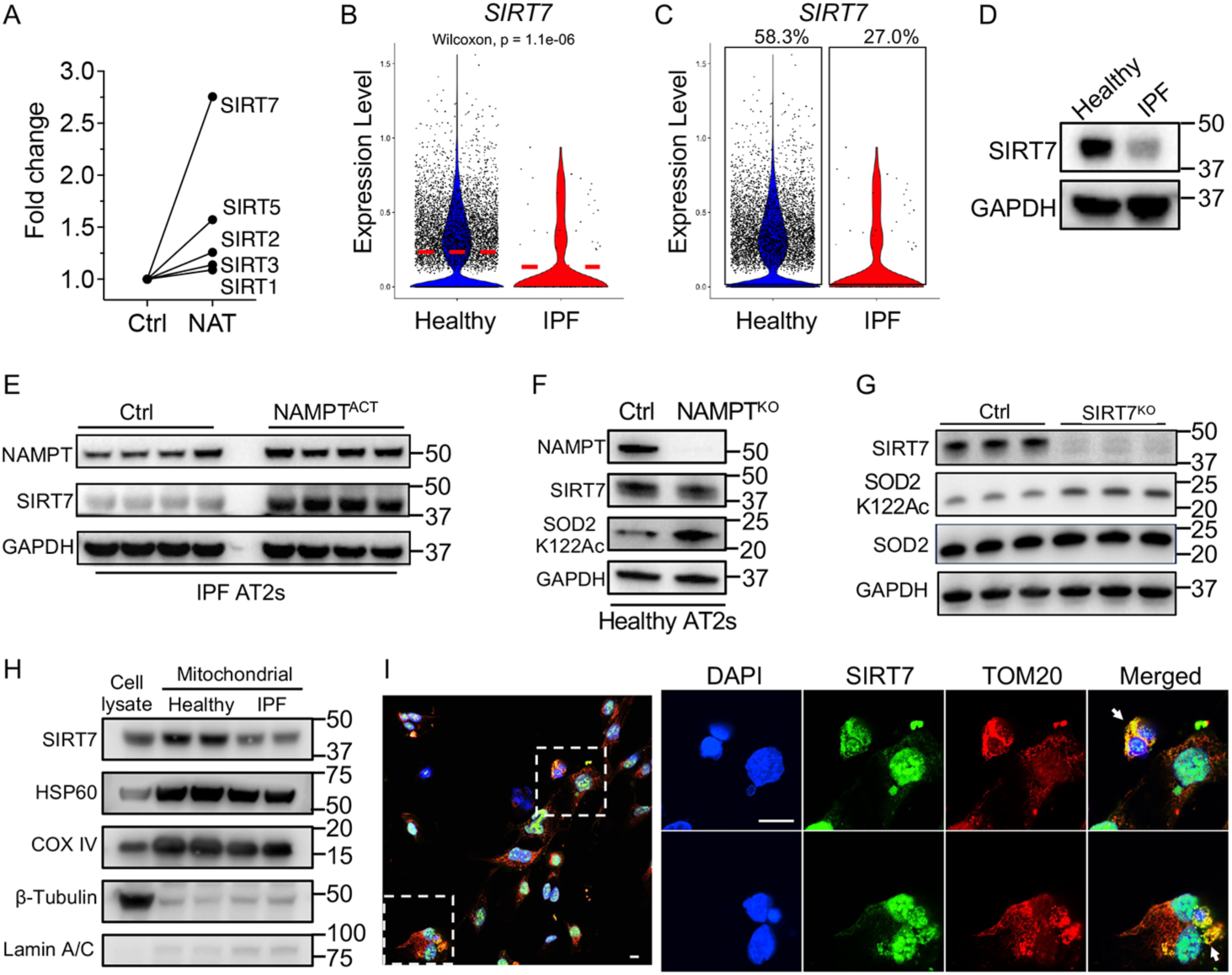
NAMPT regulates SIRT7 expression and SIRT7 deacetylates SOD2 in AT2s. (A) Fold change in the expression of sirtuin family genes in AT2s from IPF patients treated with NAT or DMSO, analyzed by bulk RNA-seq. (B and C) Violin plots showing (B) *SIRT7* expression levels and (C) the percentage of *SIRT7*^+^ cells in AT2s from healthy and IPF lungs from in house scRNA-seq dataset. (D) Western blot analysis of SIRT7 expression in immortalized AT2s from healthy and IPF lungs. GAPDH served as loading control. (E) Western blot analysis of the expression of NAMPT and SIRT7 in NAMPT^ACT^ and control immortalized AT2s. GAPDH served as loading control. (F) Western blot analysis of the expression of NAMPT, SIRT7, and SOD2 K122Ac in NAMPT^KO^ and control immortalized AT2s. GAPDH served as loading control. (G) Western blot analysis of the expression of SIRT7, SOD2 K122Ac and total SOD2 in SIRT7^KO^ and control immortalized AT2s. GAPDH served as loading control. (H) Western blot analysis of SIRT7, mitochondrial proteins (HSP60 and COX IV), nuclear protein (Lamin A/C), and cytosolic protein (β-Tubulin) in mitochondria-isolated fractions from immortalized AT2s from healthy and IPF lungs. Whole cell lysate was used as a control. (I) Immunofluorescence co-staining of SIRT7 and TOM20 in immortalized healthy AT2s (Left). Representative cells (indicated by arrows) are shown at higher magnification (Right). Scale bar, 100 μm.

SIRT7 protein is primarily localized to the nucleus^73,74^. However, previous studies demonstrated that SIRT7 is also found outside the nucleus^37,74,75^. A recent published study showed that SIRT7 protects neural stem cells by suppressing mitochondrial protein folding stress, suggesting SIRT7 functions in mitochondria^76^. We performed western blot analysis of mitochondrial proteins isolated from immortalized healthy and IPF AT2s to examine if SIRT7 was present in mitochondria. The data revealed that SIRT7 was present along with the classical mitochondrial markers, HSP60 and complex IV subunit IV (COX IV) (Figure 4H). Interestingly, the abundance of SIRT7 in the mitochondria protein isolated from IPF AT2s was lower than that from healthy AT2s (Figure 4H).

Next, we performed immunofluorescence co-staining of SIRT7 and mitochondrial protein TOM20 in cultured immortalized human AT2s. We observed cells with abundant co-expression of SIRT7and mitochondrial marker TOM20 while SIRT7 is highly presented in the nucleus (Figure 4I). Together, these data suggest that SIRT7 is also expressed in mitochondria where it might activate mitochondrial SOD2.

### NAMPT regulates mitochondrial function and AT2 cell differentiation

Next, we investigated the role of NAMPT in regulating mitochondrial function of AT2s. We observed that NAMPT^low^ AT2s had decreased mitochondria related gene expression scores^77^ compared to the NAMPT^hi^ AT2s (Figure 5A). Seahorse mitochondrial stress testing revealed that IPF AT2s had decreased oxygen consumption rate (OCR) compared to that of healthy AT2s (Figure 6B). Importantly, the NAMPT inhibitor, FK866, decreased the OCR of healthy AT2s (Figure 5C) which suggested NAMPT has a direct role in regulating mitochondrial function of AT2s. On the other hand, activating NAMPT in IPF AT2s with NAT treatment increased expression of mitochondrial related genes including *COX5B*, *PINK1*, and *MFN2* (Figure 5D). Pathway analyses showed that NAMPT activation was positively correlated with the signaling of the biocarta mitochondria pathway (Figure 5E). Functionally, NAMPT activation in AT2s increased mitochondrial membrane potential (Figure 5F), increased total ATP production (Figure 5G), and ATP production from mitochondrial respiration in AT2s (Figure 5G, red bars).

**Figure 5.**
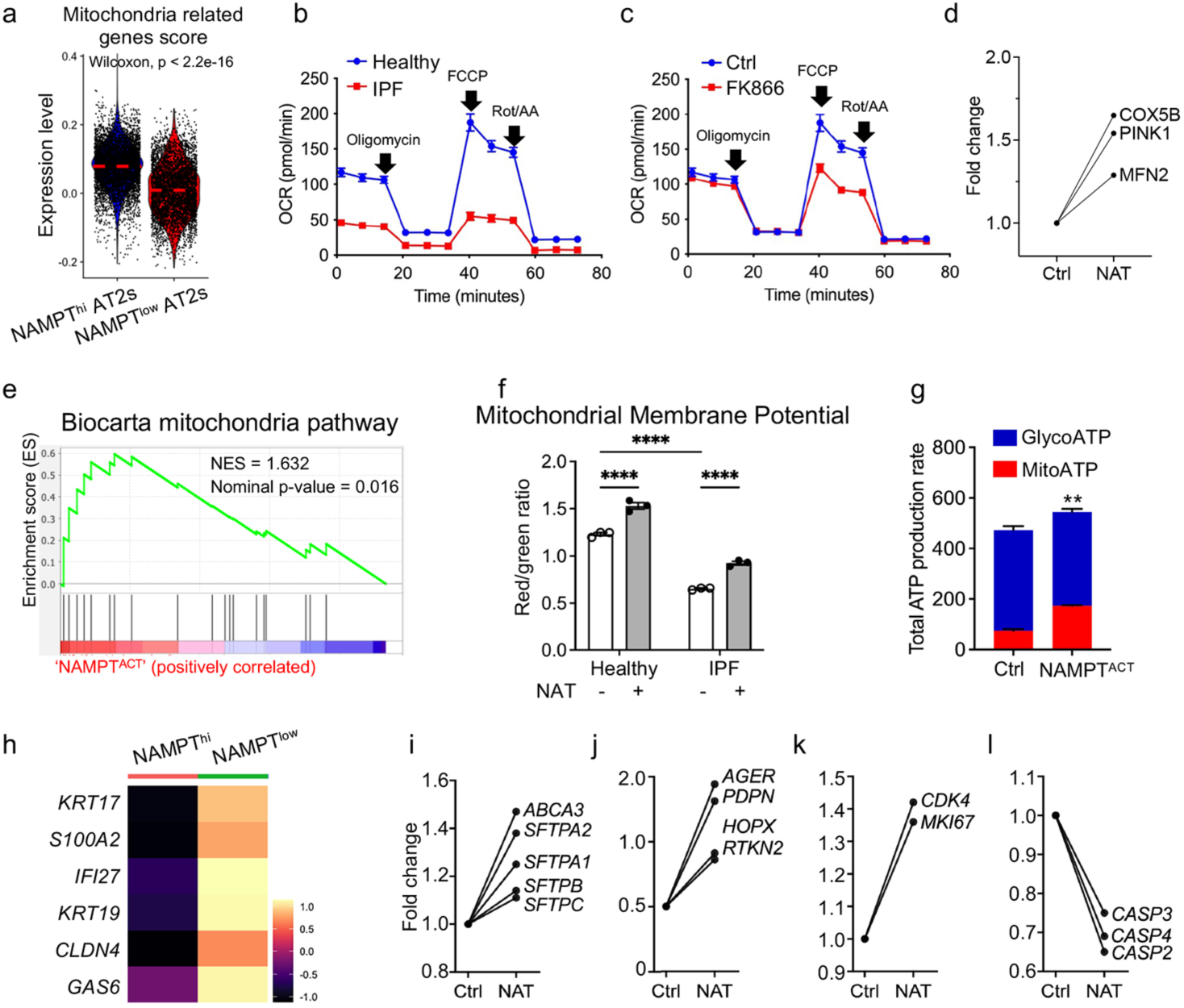
NAMPT regulates mitochondrial function and AT2 cell differentiation. (A) Violin plots comparison of the mitochondrial related gene score in NAMPT^hi^ and NAMPT^low^ human AT2s. (B and C) Oxygen consumption rate (OCR) of (B) immortalized healthy and IPF AT2s and (C) immortalized healthy AT2s treated with FK866 or vehicle control was measured by Seahorse analysis. Data are shown as the mean ± SEM from 6 replicate wells. (D) Fold change in the expression of mitochondrial genes in AT2s from IPF patients with NAT or DMSO control, analyzed by bulk RNA-seq. (E) Gene set enrichment analysis (GSEA) plots for Biocarta mitochondria pathway in NAMPT^ACT^ compared with control immortalized AT2s. Green curves indicate enrichment scores. The normalized enrichment score (NES), FWER P value are indicated within the graph. NES > 1.1 and p < 0.05 were considered statistically significant. (F) Mitochondrial membrane potential of immortalized healthy and IPF AT2s with or without NAT treatment evaluated by JC-1 staining followed by flow cytometry. (n = 3 per group, ****p < 0.0001 by ANOVA). (G) Mitochondrial ATP production of immortalized IPF AT2s with NAMPT activation by CRISPR/Cas9 and control cells measured by Seahorse XF Real-Time ATP Rate Assay kit. Data are shown as the mean ± SEM from 6 replicates. **p < 0.01 by unpaired t test. (H) Heatmap of upregulated transitional cell marker genes in NAMPT^hi^ versus NAMPT^low^ human AT2s. (I-L) Fold change in the expression of (I) AT2, (J) AT1 cell markers, (K) proliferation markers, (L) CASPASE genes in AT2s from IPF patients treated with NAT versus DMSO control, analyzed by bulk RNA-seq. Data are shown as the mean ± SEM.

**Figure 6.**
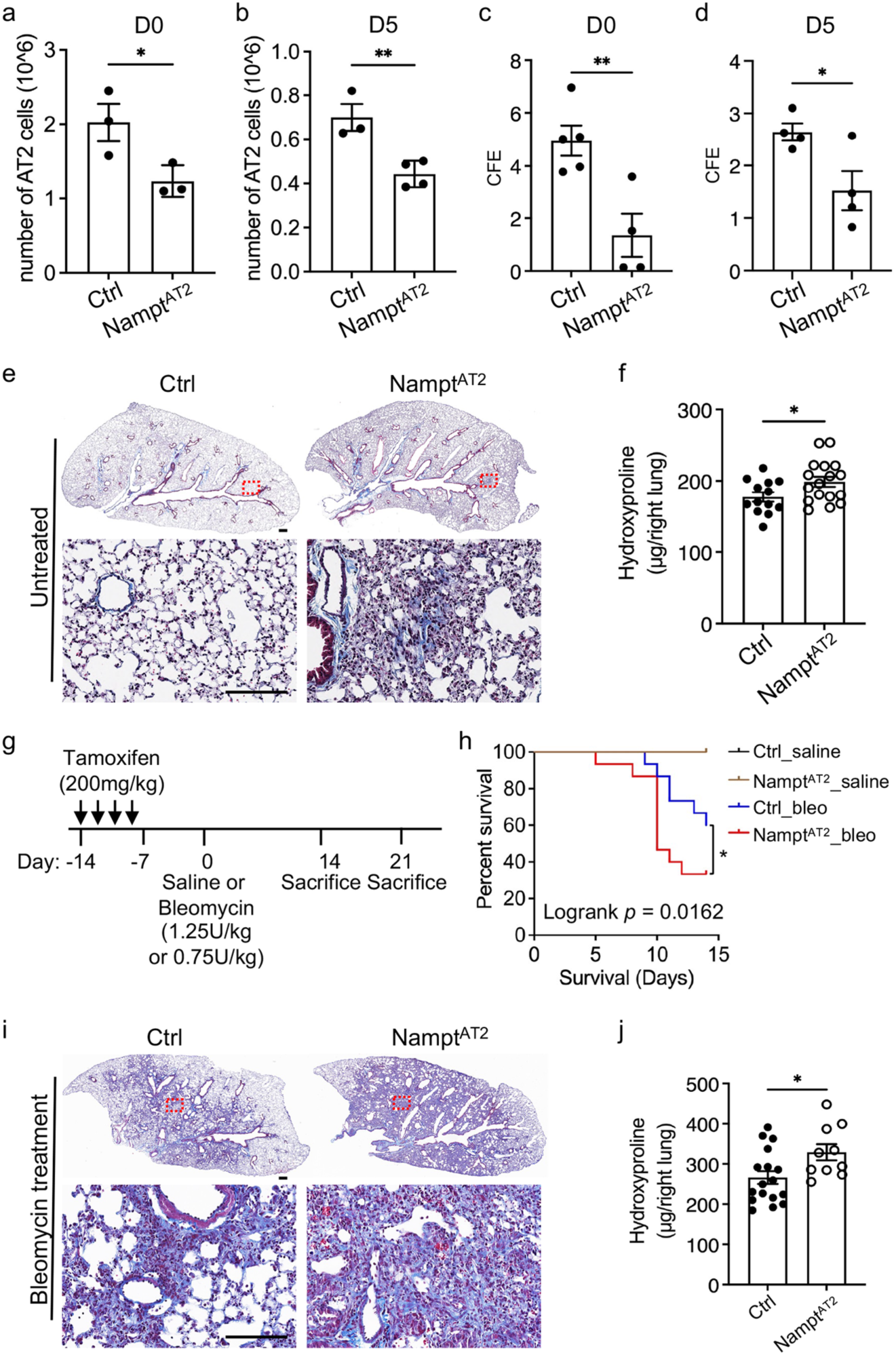
AT2 specific Nampt deletion impaired AT2 progenitor renewal in vivo and increased lung fibrosis. (A and B) Cell number of AT2s recovered per lung from (A) uninjured (D0) and (B) 5 days after bleomycin injury (D5) Nampt^AT2^ and control mice (n =3-4 per group). *p < 0.05, **p < 0.01 by unpaired t test. (C and D) CFE in 3D organoid cultures of AT2s from Nampt^AT2^ and control mice, (C) uninjured (D0) or (D) 5 days after bleomycin injury (D5) (n =4-5 per group). *p < 0.05, **p < 0.01 by unpaired t test. (E and F) Spontaneous lung fibrosis developed in Nampt^AT2^ mice at the age of 14 months. (E) Trichrome staining and (F) Hydroxyproline contents (n = 13 – 17 per group). *p < 0.05 by unpaired t test. Scale bar, 500 μm. (G-J) Experiment of young Nampt^AT2^ and control mice (8-10 weeks old) treated with bleomycin and sacrificed on day 14 or day 21. (G) Experiment layout of Nampt^AT2^ and control mice treated with bleomycin following tamoxifen injection. (H) Survival curve (n = 15, *p < 0.05 by log rank (Mantel-Cox) test) of young Nampt^AT2^ and control mice treated with 1.25 U/kg bleomycin and sacrificed on day 14. (I) Trichrome staining and (J) Hydroxyproline contents in the lungs of young Nampt^AT2^ and control mice (8-10 weeks old) treated with 0.75 U/kg bleomycin and sacrificed on day 21 (n = 13-17 per group). *p < 0.05 by unpaired t test. Scale bar, 500 μm. Data are shown as the mean ± SEM. See also Figure S3

Interestingly, NAMPT^low^ AT2s also showed increased expression of transitional cell marker genes including *KRT17*, *KRT19*, and *GLDN4* compared to NAMPT^hi^ AT2s (Figure 5H). IPF AT2s treated with NAMPT activator, NAT exhibited increased expression of both AT2 and AT1 marker genes (Figure 5I and 5J, respectively), increased *CDK4* and *MKI67* expression (Figure 5K), and decreased expression of CASPASE genes (Figure 5L). These data suggest that NAMPT activation promotes AT2 cell proliferation and differentiation to AT1 cells as well as suppression of AT2 cell apoptosis.

### AT2-specific Nampt deletion impaired AT2 progenitor renewal in vivo and increased lung fibrosis

To examine the role of NAMPT in regulating AT2 progenitor cell function and lung fibrosis *in vivo*, we generated a novel mouse model with AT2 specific Nampt deletion (Nampt^AT2^). These mice were generated by cross-breeding the Nampt floxed mice^78,79^ with *Sftpc-CreER* mice (Figure S3A) followed by tamoxifen treatment. Nampt expression is abrogated in AT2s upon tamoxifen treatment (Figure S3B). Nampt^AT2^ mice are fertile with normal litter size and developed normally to adulthood. There were no physical abnormalities in adult Nampt^AT2^ mice (8-10 weeks old) under homeostatic conditions. scRNA-seq analysis showed that AT2s from tamoxifen treated Nampt^AT2^ mice had decreased expression of *Sirt7*, *Sod2*, and *Romo1* (Figure S3C). Nampt-deficient AT2s also showed decreased mitochondria related gene expression scores (Figure S3D). These data further confirmed the role of NAMPT in regulating oxidative stress and mitochondrial function of AT2s. Next, we compared AT2 cell recovery from tamoxifen treated Nampt^AT2^ and control mice with and without bleomycin treatment. Nampt^AT2^ mice had fewer AT2s recovered per lung compared to that of control mice both under homeostasis (Figure 6A) and on day 5 after bleomycin injury (Figure 6B). Using flow cytometry analysis, we showed that AT2s from Nampt^AT2^ mice had increased mitochondrial superoxide levels compared to that of AT2s from control mice (Figure S3E). AT2s isolated from Nampt^AT2^ mice (both uninjured - D0 and day 5 bleomycin injured - D5) displayed significantly reduced colony formation capacity compared to AT2s isolated from control mice (Figure 6C, 6D and S3F).

Nampt^AT2^ mice developed spontaneous lung fibrosis at the age of 14 months as shown by both increased trichrome staining (Figure 6E) and increased hydroxyproline contents in the lungs (Figure 6F). We then subjected the mice to bleomycin lung injury. With initial experiment, we observed that Nampt^AT2^ mice were more susceptible to bleomycin injury. We treated Nampt^AT2^ and control mice with bleomycin (1.25 U/kg) two weeks after four days of tamoxifen treatment (Figure 6G). The Nampt^AT2^ mice showed increased mortality on day 14 after bleomycin treatment (Figure 6H). We next reduced the bleomycin dose to 0.75 U/kg to decrease mortality and observed a higher number of surviving mice on day 21 for lung fibrosis assessment (Figure 6G). Nampt^AT2^ mice showed increased lung fibrosis after bleomycin lung injury compared to bleomycin-treated control mice as measured by trichrome staining (Figure 6I) and increased hydroxyproline contents (Figure 6J) in the lungs.

These observations revealed that NAMPT deficiency impaired AT2 regeneration, exacerbated bleomycin-induced fibrosis in young mice, and led to spontaneous fibrosis in aged mice, underscoring the essential role of NAMPT in alveolar repair and long-term lung homeostasis.

### NAMPT activation in vivo promoted AT2 regeneration and attenuated lung fibrosis

Wild-type mice (C57BL/6J mice; 10 weeks old) were administered NAT to investigate the potential therapeutic effect of NAMPT activation on lung fibrosis. The first treatment paradigm (Figure 7A) involved dosing of NAT (20 mg/kg) or vehicle via intraperitoneal (i.p.) injection daily from 3 days before (day −3) to 3 days after (day 3) bleomycin treatment (2.5 U/kg; day 0) (Figure 7A). Mice were sacrificed on day 5 after bleomycin treatment and lung epithelial cells were isolated and analyzed by flow cytometry and cell counting. Mice treated with both bleomycin and NAT had an increased percentage (Figure 7B and 7C) and number (Figure 7D) of total lung epithelial cells in the lungs compared to mice treated with bleomycin alone. Within the lung epithelial cells, NAT treated mice showed an increased percentage and number of AT2s recovered per lung (Figure 7B, 7E and 7F). Interestingly, NAT treatment alone did not alter the number of total BAL cells and BAL-localized macrophages on day 5 after bleomycin injury (Figure S4A and S4B), indicating that NAT treatment did not promote lung inflammation.

**Figure 7.**
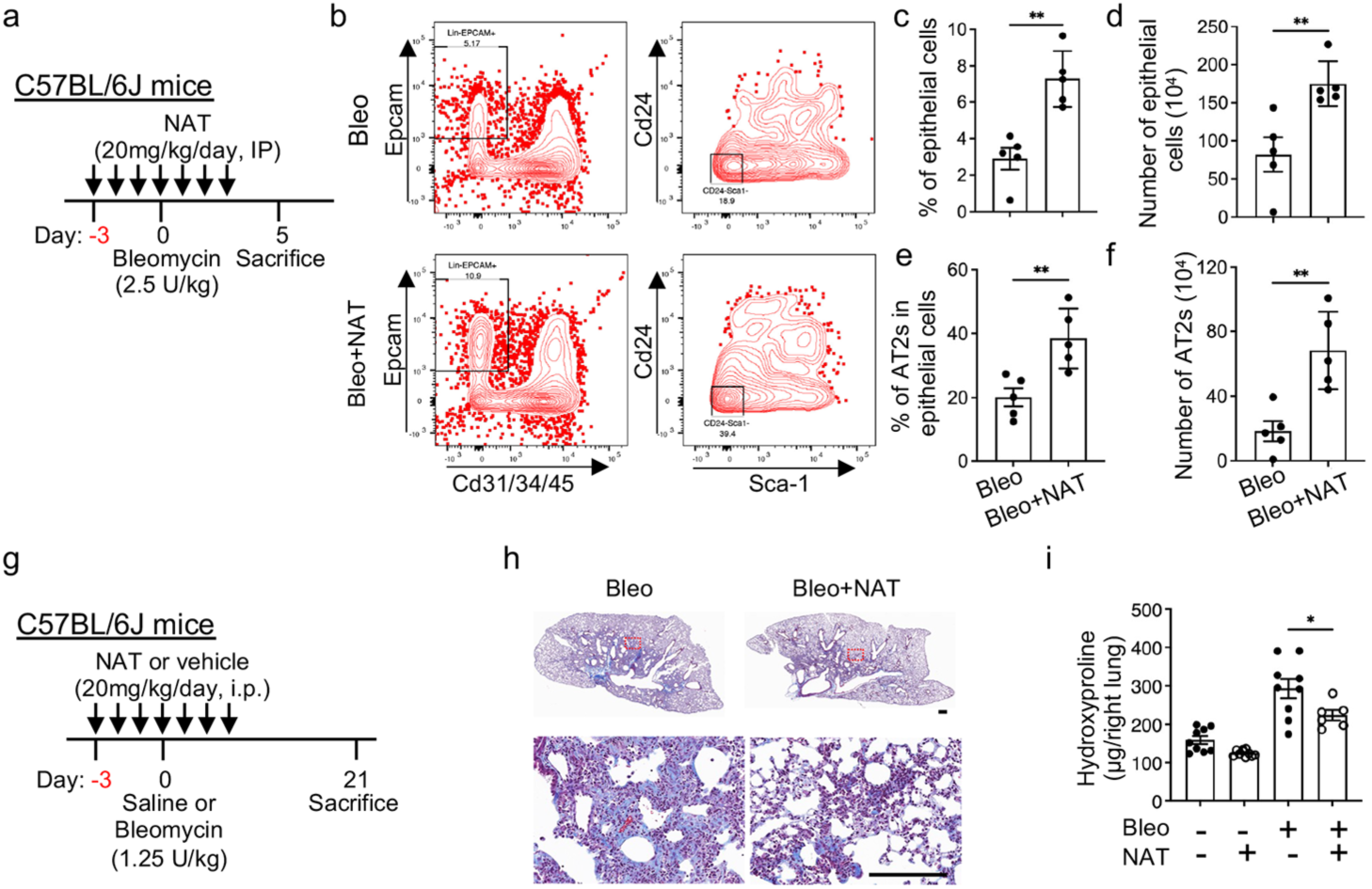
NAMPT activation in vivo promoted AT2 regeneration and attenuated lung fibrosis. (A-F) Effect of NAT treatment on AT2 cell recovery. (A) Experiment layout: Young C57BL/6J (10 weeks old) wild type mice were treated with 20 mg/kg NAT and 2.5 U/kg bleomycin and lungs were collected on day 5 after bleomycin treatment. (B) Flow cytometry gating strategies for total lung epithelial cells and AT2s. Lung epithelial cells were gated as Epcam^+^Cd31^−^Cd34^−^Cd45^−^ cells in live cell populations. AT2 were gated as CD24^−^Sca-1^−^ in epithelial cell populations. (C and D) Percentages of epithelial cells in total cells and numbers of epithelial cells per lung (n = 5 per group). **p < 0.01 by unpaired t test. (E and F) Percentages of AT2s in epithelial cells and numbers of AT2s per lung (n = 5 per group). **p < 0.01 by unpaired t test. (G-I) The prophylactic regimen of NAT treatment. (G) Experiment layout: Young C57BL/6J (10 weeks old) wild type mice were treated with 20 mg/kg NAT and 1.25 U/kg bleomycin and lungs were collected on day 21 after bleomycin treatment. (H) Trichrome staining (scale bar, 500 μm) and (I) hydroxyproline contents (n = 6-10 per group, *p < 0.05 by ANOVA) were used to assess collagen levels in mouse lungs. Data are shown as the mean ± SEM. See also Figure S4

## Discussion

Multiple studies from our group and others point to AT2 progenitor cell failure as a critical event in IPF^6,8,16,19,20^, but the molecular mechanisms that regulate AT2 progenitor renewal in lung fibrosis are not completely understood. Our discovery of a significant reduction of NAMPT in IPF AT2 progenitor cells led us to hypothesize that NAMPT deficiency in IPF AT2s may increase susceptibility of AT2s to oxidative stress and mitochondrial dysfunction, resulting in impaired AT2 cell renewal and severe lung fibrosis. NAD^+^ is a central regulator of cell energy metabolism^18,19^. NAD^+^ levels were decreased in IPF lungs^19,20^, and NAD^+^ deficiency is associated with lung fibrosis^19–21^. However, the causes of NAD^+^ deficiency in AT2s from fibrotic lungs is yet to be elucidated. In the current study, we have determined that NAMPT activity is crucial for maintaining NAD^+^ levels in AT2s, and for AT2 progenitor renewal. Inhibition of NAMPT suppressed and activation of NAMPT by small molecule activators increased renewal of both healthy and IPF AT2s in 3D organoid cultures. We further dissected the mechanisms under which NAMPT regulates oxidative stress and mitochondrial function. NAMPT deficiency in IPF AT2s downregulates SIRT7 and SOD2 which leads to increased susceptibility of AT2s to oxidative stress and mitochondrial dysfunction, resulting in impaired AT2 cell renewal and severe lung fibrosis. Most importantly, restoration of NAMPT activity by treatment with small molecule NAMPT activators increased AT2 recovery and attenuated bleomycin induced lung fibrosis in vivo with both young and old mice. This work provides mechanistic evidence and highlights potential avenues for the development of novel therapies for severe pulmonary diseases such as IPF.

Expression of oxidative stress response proteins have been known to be reduced in IPF epithelial cells^52–54^ and with the ensuing loss of cellular redox homeostasis has been suggested to be a fundamental driver of IPF pathogenesis for decades^50,80^. In fact, SOD2 was shown to be reduced in IPF BAL which prompted clinical investigations in IPF patients aimed at restoring antioxidant function^81^. In the current study, we have found that NAMPT deficiency is associated with downregulation of SOD2 in AT2s which results in their increased susceptibility to oxidative stress and impaired progenitor cell renewal. This provides a critical novel missing link identifying the underlying defect in oxidative stress responses in IPF as being related to NAD+ metabolism.

We have previously reported that downregulation of NAD^+^-dependent sirtuin signaling contributed to the impaired progenitor function of IPF AT2s, and NAD^+^ precursors promoted AT2 renewal^8^. Our current results show that NAMPT activation dramatically upregulated SIRT7 expression while the expression of other sirtuin family member genes were increased moderately. Knockout of NAMPT in human AT2s decreased SIRT7 expression and decreased SOD2 activation. Furthermore, direct knockout of SIRT7 in AT2s resulted in increased K122-SOD suggesting that SIRT7 deacetylates SOD2 in AT2s. SIRT7 primarily resides in the nucleus^73,74^. However, the presence of SIRT7 in extranuclear locations has been demonstrated previously^37,74,75^. In this study, we provide evidence by both western blot and immunofluorescence staining showing that SIRT7 is also present in mitochondria in AT2s. Furthermore, we showed that SIRT7 deacetylates and actives SOD2 in AT2s. All sirtuins are NAD^+^ dependent. We do not rule out that other sirtuin members may also participate in NAMPT activation associated SOD2 activation even though we have found that NAMPT activation upregulated SIRT7 most significantly. For example, SIRT3 has been identified as a deacetylase for SOD2 in lung epithelial cells and other cell types^56,71^.

The fundamental role of the lung is to facilitate gas exchange in the distal alvealor space and this requires the generation of oxygen transmitting AT1 cells from the surfactant producing progenitor AT2 cells. IPF patient succumb to hypoxemia becasue of the loss of AT2 and AT1 cells and the inability to transfer oxygen from ambient air into blood. Impaired type II to type I alveolar epithelial cell transition and accumulation of pathologic transitional cells in fibrotic lungs has been identified in the lungs of both IPF patients and mice with experimental lung fibrosis by multiple reports but the mechanism has been unknown^62,82–84^. Interestingly, we have observed increased expression of transitional cell marker genes in NAMPT^low^ human AT2s, suggesting that NAMPT deficiency may also impair AT2 cell to AT1 cell differentiation. The role of NAMPT in regulating AT2 cell differentiation is one of the focuses of further studies in the lab. Existing therapies to treat IPF target fibroblast production of collagen and slow the loss of lung function but patients do no improve functional status, demonstrate improved symptoms and have not prospectively been shown to have improved mortality. This is likely because these therapies do not target the underlying cause of IPF which is AT2 failure. IPF AT2 cells have been shown to have short telomeres and this likely contributes to failed regenerative capacity. Short telomeres have been shown to correlated with mortality^85^. Studies are ongoing to determine if there is a link between NAMPT deficiency and telomere shortening.

Extracellular NAMPT (eNAMPT) functions as a damage-associated molecular pattern (DAMP) protein that regulates inflammation in acute lung injury through endothelial cells and macrophages^79,86–96^. In our studies, we did not observe increased number of macrophages in BAL from NAMPT activator, NAT, treated mice.

NAMPT is also expressed in fibroblasts. Published study showed NAD+ metabolism plays a role in regulating lung fibroblasts phenotype and apoptosis^97^. Our scRNA-seq analysis showed that NAMPT expression levels were not significantly changed in fibroblasts from bleomycin injured mouse lungs and IPF lungs. More studies on the role of NAMPT in the interactions between AT2s and macrophages, and between AT2s and fibroblasts in lung fibrosis is needed in the future.

In this study, we discovered that IPF AT2s have reduced levels of cellular NAMPT. NAMPT deficiency downregulates SIRT7 and SOD2 expression which results in increased oxidative stress, mitochondrial dysfunction, and impaired AT2 progenitor renewal in IPF. This observation was made from IPF explants where patients have essentially succumbed to respiratory failure. This observation from IPF patient highlights the importance of the observation that was then used to develop a novel mouse mode of IPF. Our novel mouse model (Nampt^AT2^) with AT2 specific NAMPT deletion had decreased AT2 renewal and reduced recovery from the lungs. AT2s from the Nampt^AT2^ mice exhibited decreased SIRT7 and SOD2 expression and increased mitochondrial superoxide levels. The Nampt^AT2^ mice exhibited worse survival and increased lung fibrosis after bleomycin injury and developed spontaneous lung fibrosis at older age. These data indicated that our mouse model recapitulates the key aspects of severe pulmonary fibrosis in human disease. Excitingly, we have shown that small molecule NAMPT activators not only promoted AT2 renewal in 3D organoid culture and restored AT2 homeostasis, but also attenuated bleomycin induced lung fibrosis in vivo with both prophylactic and therapeutic benefits. These results suggest we have discovered the “Achilles Heel” of IPF and provide novel insights for developing novel therapeutics for the fatal disease IPF that target the underlying cause and not the downstream effects and most importantly offer hope to improve form, function, and lifespan.

### Limitations of the study

One limitation of our study is the technical difficulty in flow-sorting sufficient numbers of AT2 progenitor cells to measure NAD⁺ levels, due to their reduced abundance in IPF lungs. To address this, we successfully generated immortalized human AT2 cells from both healthy and IPF lungs, and used these cells to assess the effects of NAT treatment on NAD⁺ biosynthesis. As an alternative approach, we plan to use iPSC-derived human AT2 cells (BU3-NGST) obtained from Dr. Darrell Kotton^98^.

Another limitation is that we did not examine the role of extracellular NAMPT (eNAMPT) in our study. Prior reports have shown that eNAMPT promotes inflammation in acute lung injury through effects on endothelial cells and macrophages^79,86–96^. Whether NAT treatment influences eNAMPT levels or inflammatory responses in the bleomycin-induced lung fibrosis model remains unknown. Although our initial experiments indicated that NAT treatment enhanced AT2 recovery (Figure 7B-7I), we observed no significant changes in lung inflammation (Figure S4A and S4B). We will continue to monitor inflammatory responses in NAT-treated mice, and plan to investigate eNAMPT expression further if inflammation becomes evident. Additionally, we aim to measure circulating eNAMPT levels in blood samples from IPF patients in future studies.

## RESOURCE AVAILABILITY

### Lead Contact

Further information and requests for resources and reagents should be directed to and will be fulfilled by the Lead Contact, Carol Jiurong Liang.

### Materials Availability

All unique/stable reagents generated in this study are available from the Lead Contact with a completed materials transfer agreement.

### Data and Code Availability

The raw data files of the single cell RNA-seq are deposited as GSE295739 and GSE295740. Passcodes for reviewers are anunowyetdaxjgd and cfqnaeacnledrqv, respectively. R code files used for data integration and analysis are available at https://github.com/jiang-fibrosis-lab. Other scRNA-seq data used in this paper can be accessed via datasets GSE146981^64^, GSE128033^61^, GSE135893^63^, GSE136831^62^, GSE122960^99^ and GSE132771^100^.

## ACKNOWLEDGMENTS

We thank the members of our laboratory for support and helpful discussion during the course of the study. This work was supported by grants from National Institutes of Health to J.L. (R01 AG078655), P.W.N. (R35-HL150829 and P01-HL108793), D.J. (R01 HL172990), and awards to X.L. from American Heart Association (Career Development Award 24CDA1268568) and Pulmonary Fibrosis Foundation (PFF Scholars Program 1272558).

## AUTHOR CONTRIBUTIONS

J.L., P.W.N., and D.J. conceived the study. X.Z. performed most of the experiments and analyzed the data. X.L. analyzed single cell RNA transcriptome data, performed experiments. Y.Q., A.R., and N.L. took part in mouse, cell culture, and biological experiments. C.Y. analyzed single cell RNA transcriptome data. T.P. and P.C. provided human samples and interpreted data. P.C. and B.S. interpreted data and contributed with comments on the manuscript. X.Z., J.L., D.J., and P.W.N. wrote the paper. All authors read and reviewed the manuscript.

## STAR METHODS

### KEY RESOURCE TABLE

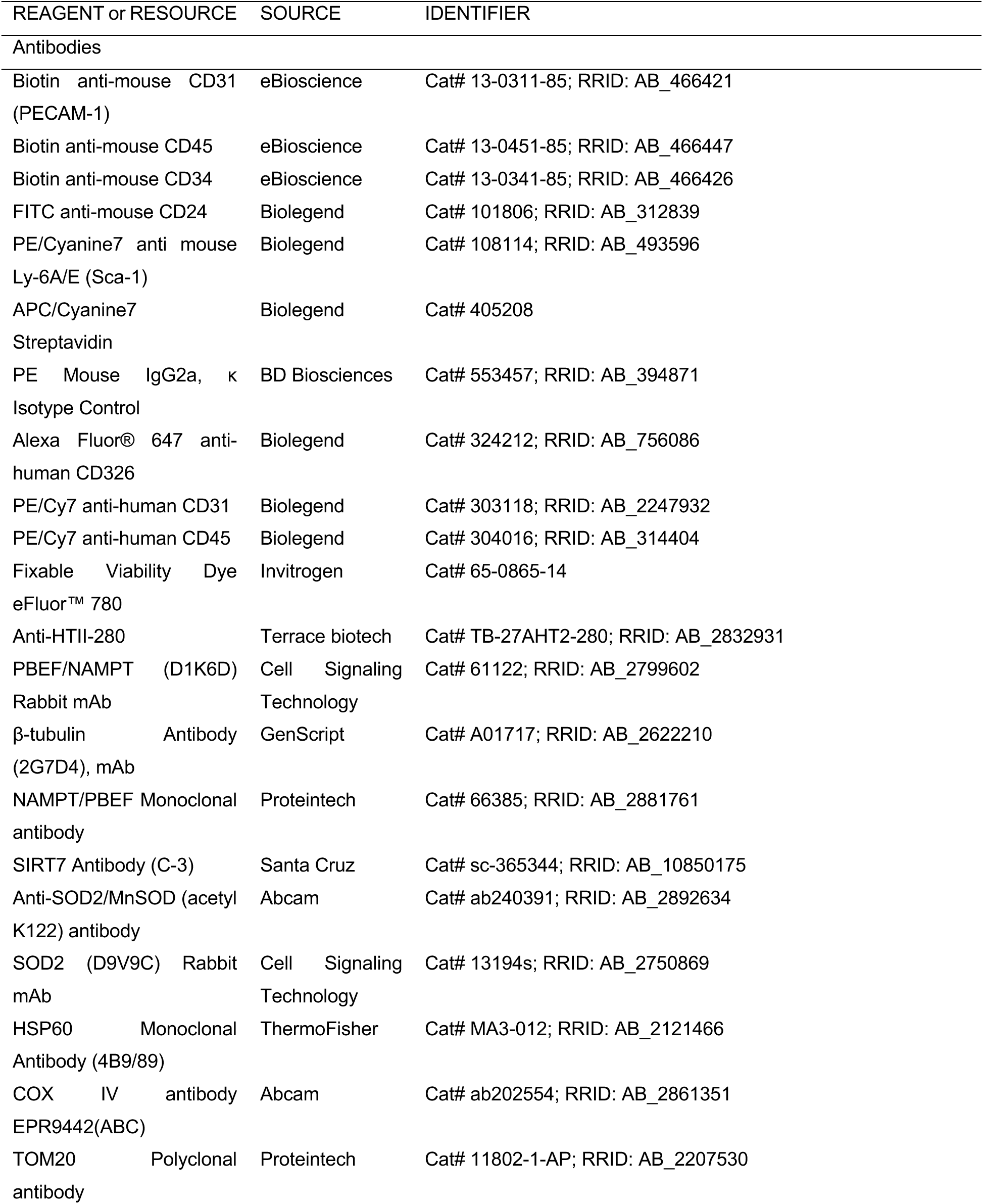
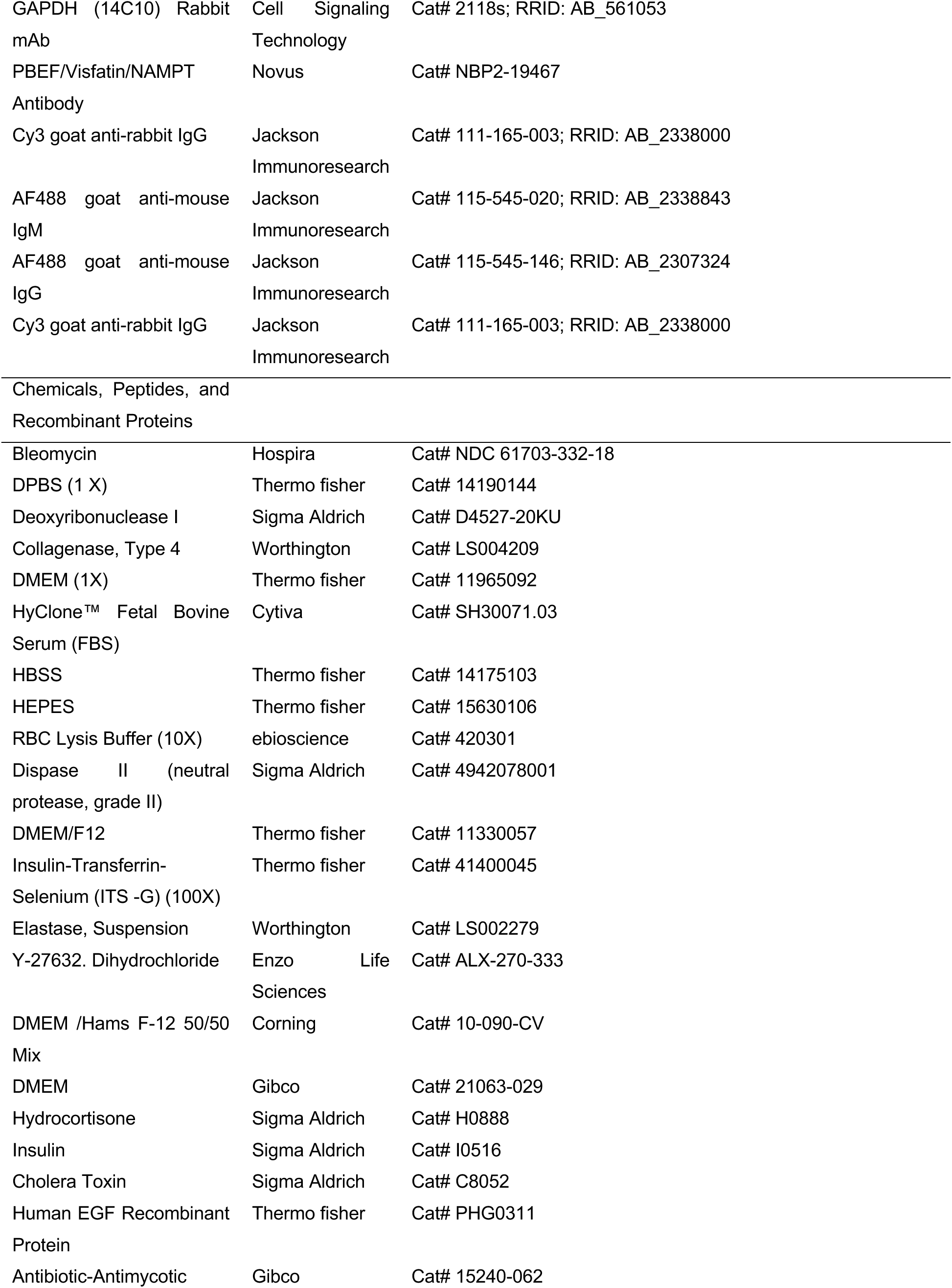
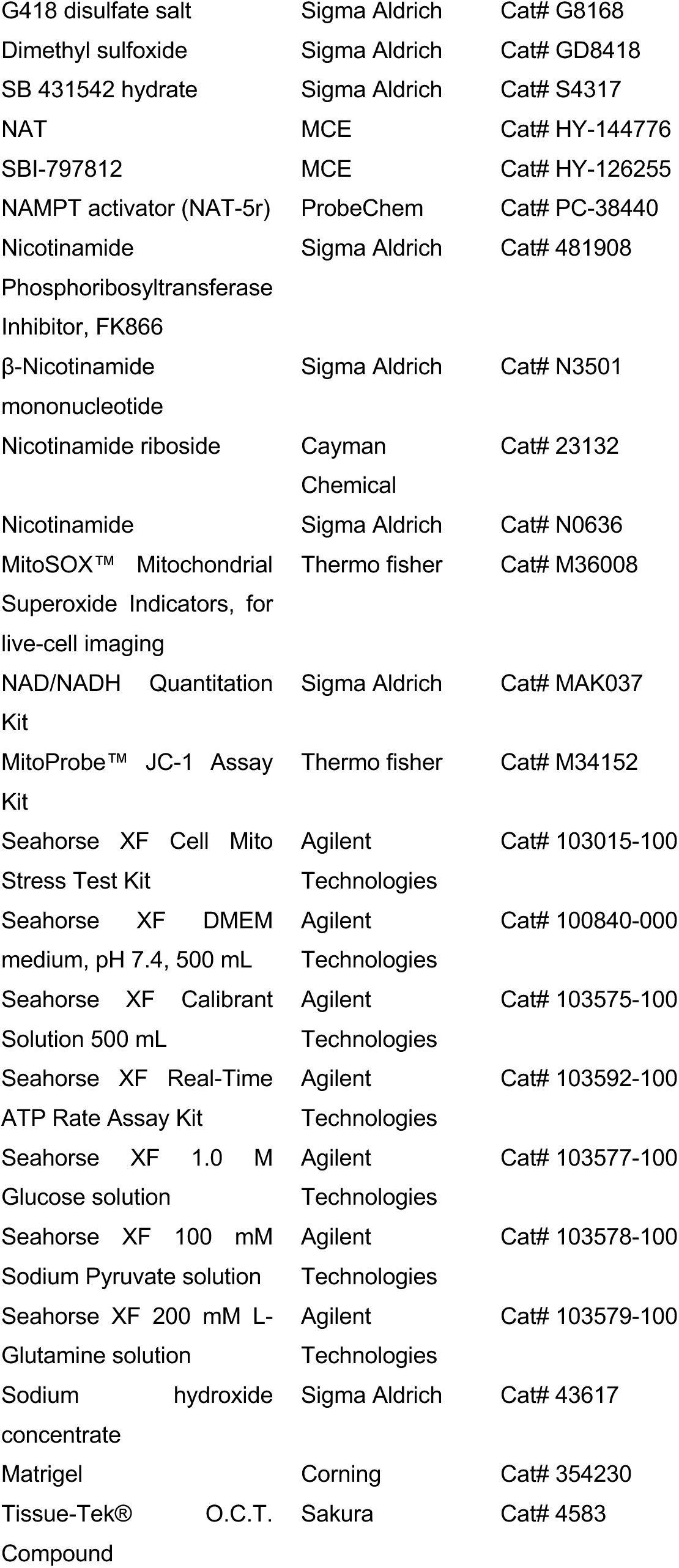
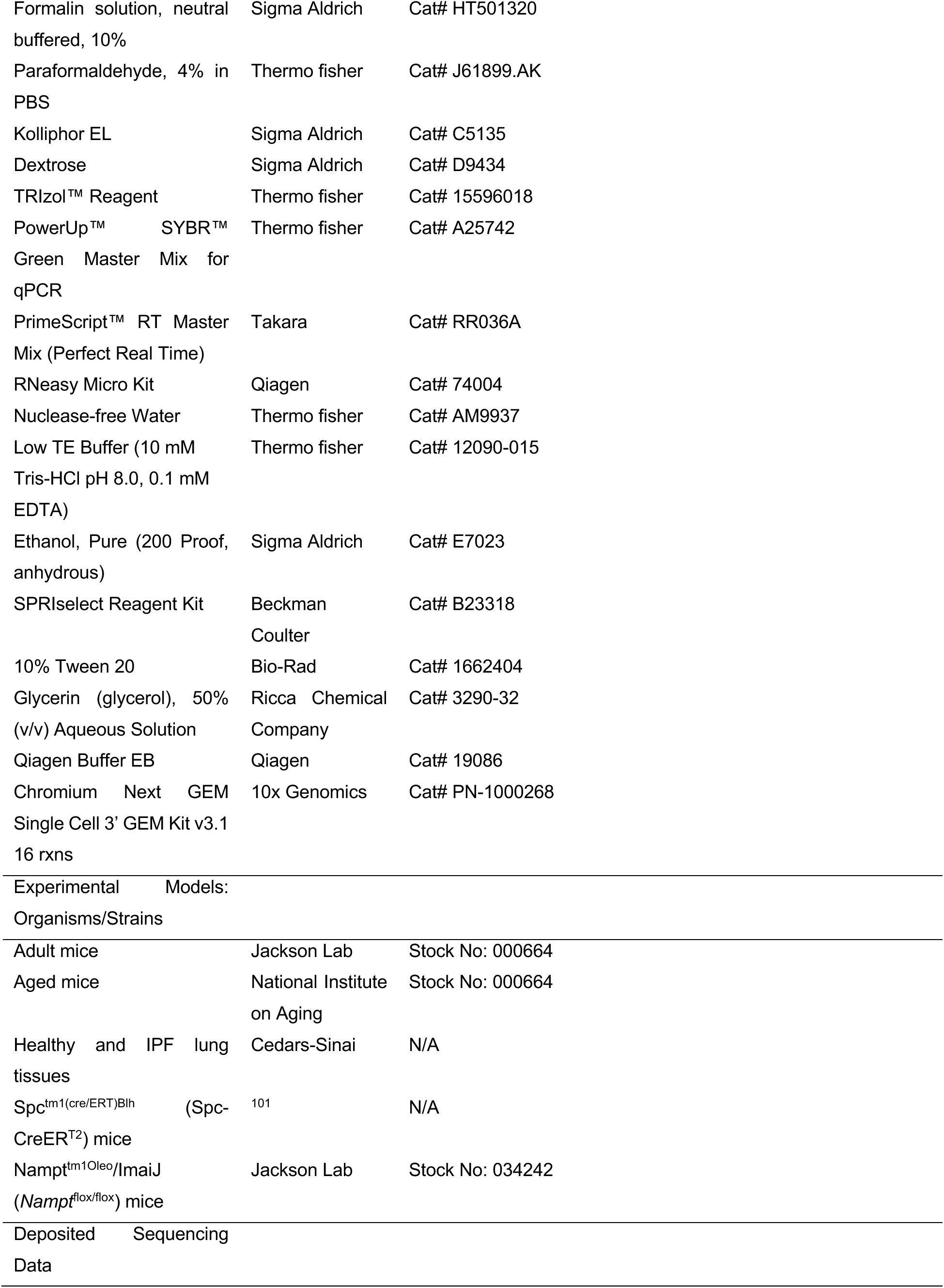
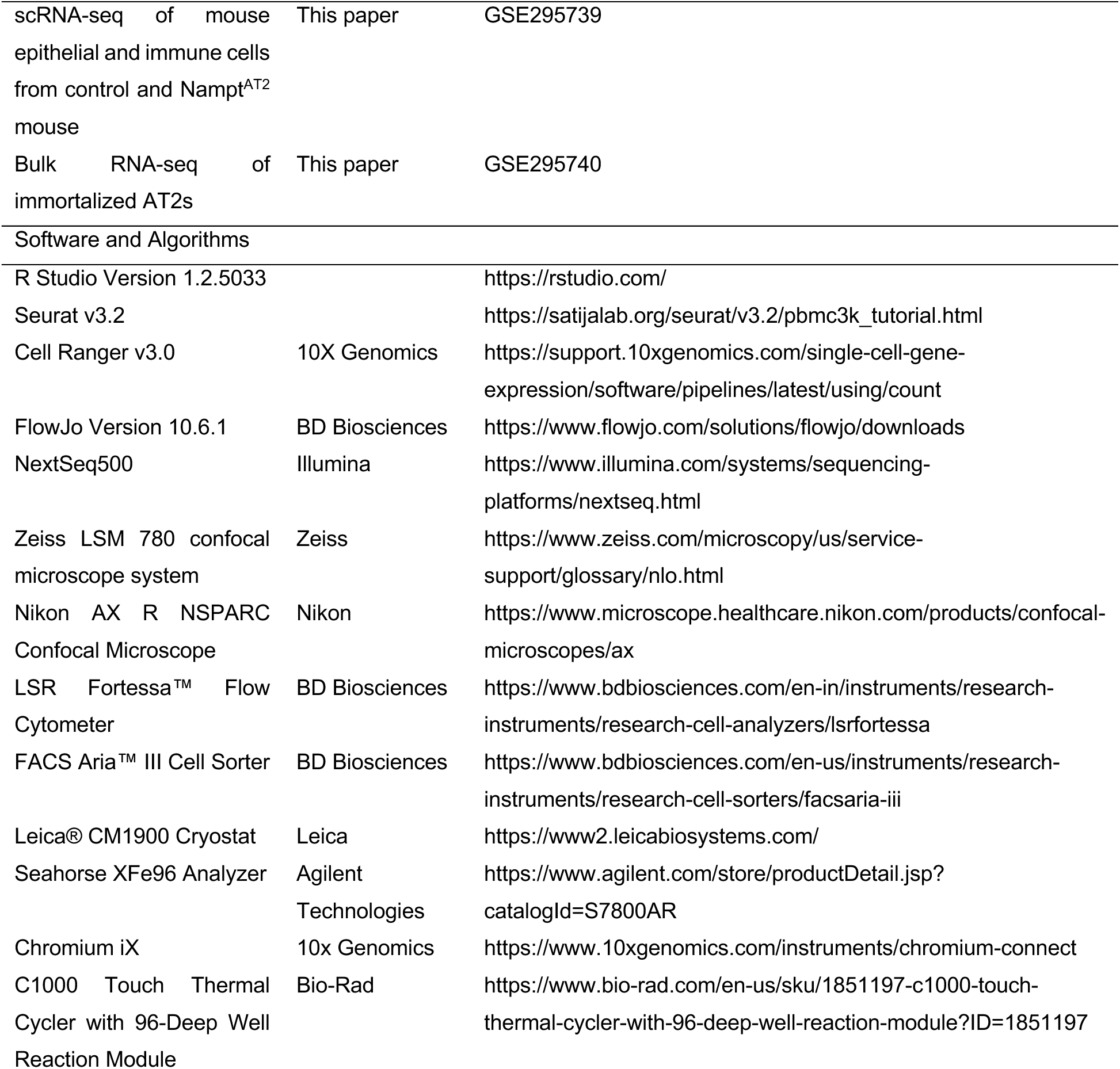

### EXPERIMENTAL MODEL AND STUDY PARTICIPANT DETAILS

#### Study approvals

Human tissue use for this study was approved by the Institutional Review Boards of Cedars-Sinai Medical Center (Pro00035409) with informed consent obtained from each participant. The study included 15 healthy donors and 18 IPF patients, comprising both males and females. All animal experiments were approved by the Institutional Animal Care and Use Committee at Cedars-Sinai Medical Center (Protocol IACUC008529). All mice were housed in a pathogen-free facility at Cedars-Sinai Medical Center and had access to auto-cleaved water and pelleted mouse diet ad libitum.

#### Mice

*SFTPC-CreER* mice was described previously^13^. Nampt floxed mice was purchase from Jackson Lab (Strain #:034242, RRID: IMSR_JAX:034242)^78^. Nampt^AT2^mice were generated by crossbreeding *SFTPC-CreER* mice and Nampt floxed mice. All mice have been backcrossed on C57BL/6J background more than six generations. Young wild-type C57BL/6J mice (8-10 weeks old) were obtained from The Jackson Laboratory and housed in the institution facility at least 2 weeks before experiments. 20-24 mon old wild-type C57BL/6J mice were obtained from National Institute on Aging. Animals were randomly assigned to treatment groups, and they were age- and sex-matched.

### METHOD DETAILS

#### Bleomycin instillation and bronchoalveolar lavage

Intratracheal bleomycin instillation was performed by following the standard prodedure described previously^8,102^. Detailed methods can be found in Supplementary Methods.

#### Human and mouse lung tissue dissociation

Human lung single cell isolation and flow cytometer were performed as previously described^6^. In brief, human lung tissues were minced and then digested with 2 mg/ml Dispase II, followed by 10 U/ml elastase and 100 U/ml DNase I digestion. Finally, cells were filtered through 100 μm cell strainer and lysed with red blood cell lysis to get single cell suspension. Detailed methods can be found in Supplementary Methods.

#### *In vitro* culture of immortalized human AT2s

Immortalized transformed AT2 cell lines were modified from Dr. Offringa group’s protocol^103^. Briefly, isolated healthy and IPF AT2s were resuspended in Fmed + ROCKinh medium, consisting of a 3:1 ratio of DMEM/F12 (Corning, Catalog 10-090-CV) to DMEM (Gibco, Catalog 21063-029), supplemented with 5% FBS, 0.4 mg/mL hydrocortisone (Sigma-Aldrich, Catalog H0888), 5 mg/mL insulin (Sigma-Aldrich, Catalog I0516), 8.4 ng/mL cholera toxin (Sigma-Aldrich, Catalog C8052), 10 ng/mL human recombinant EGF (ThermoFisher, Catalog PHG0311), Antibiotic-Antimycotic (Gibco, Catalog 15240-062), and 10 mM Y-27632 (Enzo Life Sciences, Catalog 270–333). The cells were allowed to attach for two days, with media changes every 2–3 days, and monitored daily for survival and proliferation. Once AT2s reached passage 3–4, they were infected with SV40 LgT lentivirus (customized from VectorBuilder) for immortalization, followed by G418 (Sigma-Aldrich, Catalog G8168) antibiotic selection. Single clones were sorted using HTII-280 in 96-well plates, and all AT2s were maintained in Fmed + ROCKinh medium.

#### 3D organoid cultures of human and mouse AT2s

Flow sorted AT2 cells were cultured in matrigel/medium mixture in the presence of MLg2908 lung fibroblasts^6,8^. Detailed methods can be found in Supplementary Methods.

#### Generation of NAMPT-KO, NAMPT-activation, SIRT7-KO cell line with CRISPR system

For knock out, immortalized AT2s were infected with commercial NAMPT sgRNA CRISPE/Cas9 All-in-One Lentivirus (Abm, Catalog #31414111) or commercial SIRT7 sgRNA CRISPE/Cas9 All-in-One Lentivirus (Abm, Catalog # 43812111). For activation, Immortalized AT2s were infected with commercial NAMPT CRISPRa sgRNA lentivirus (Abm, Catalog # 31414121). The efficiency of knock out or activation were validated by western blot.

#### Seahorse metabolic flux measurements

The Seahorse XF Cell Mito Stress Test (Agilent Technologies, 103015-100) and Seahorse XF Real-Time ATP Rate Assay Kit (Agilent Technologies, 103592-100) were used to assess oxygen consumption rate (OCR) and measure total ATP production rates in living cells, respectively, following the manufacturer’s instructions. Briefly, 1 × 10^4^ cells were seeded in Seahorse XF microplates. One hour before analysis, the culture medium was replaced with Seahorse XF assay media (Agilent Technologies, 103575-100), and plates were incubated at 37°C in a CO₂-free incubator. The mitochondrial stress test was conducted under basal conditions and following sequential injections of 1.5 μM oligomycin, 0.5 μM FCCP, and 0.5 μM rotenone/antimycin A. The Real-Time ATP Rate Assay was performed under basal conditions and after sequential injections of 1.5 μM oligomycin, and 0.5 μM rotenone/antimycin A.

#### Mitochondrial ROS measurement

Mitochondrial ROS production was assessed using the red MitoSox Superoxide indicator (Thermo Fisher Scientific, M36008), which detects ROS content specifically targeted to mitochondria. MitoSox Red reagent was dissolved in vehicle to make 5mM stock solution, and 5 μM working solution was prepared in HBSS. vehicle or NAT-treated immortalized AT2s were incubated with 1 mL MitoSox working solution for 20 min and washed 3 times with warm PBS. Fluorescence signals were determined by flow cytometry. To validate the MitoSOox staining results, cells were treated with DMSO as a negative control and 50 μM CCCP as a positive control.

#### NAD/NADH measurement

Total NAD and NADH were measured by NAD/NADH Quantification Kit (Sigma, Catalog MAK037) following manufacturer’s instruction. 2 × 10^5^ immortalized AT2s were washed with cold PBS, resuspended in 400 μl of NAD/NADH extraction buffer by freeze/thawing for 2 cycles of 20 minutes on dry ice followed by 10 minutes at room temperature. The extracted supernatant was split in half for total NAD^+^ and NADH after centrifugation at 13,000 × g. One-half sample was heated to 60 °C for 30 min to detect NADH. For total NAD measurement, 50 μl of sample was transferred into 96-well plate, and added 100 μl of master reaction mix (NAD cycling buffer and enzyme mix from the kit), incubated for 5 min at room temperature to convert NAD to NADH. 10 μl of developer reagent was added to each well, and incubated at room temperature for 1 h. Absorbance at 450 nm was determined using a microplate reader. and their ratios were calculated by the following equation: ratio = (NAD_total_ – NADH)/ NADH.

#### Mitochondrial membrane potential measurement

Mitochondrial membrane potential (MMP) was assessed in control and NAT-treated AT2s using the JC-1 probe (ThermoFisher, Catalog M34152). Cells were stained with 2 μM JC-1 following a previously established protocol^104^ and analyzed via flow cytometry. To validate the JC-1 results, 50 μM CCCP staining was also performed.

#### ScRNA sequencing data analysis and data collection

The human lung epithelial scRNA-seq dataset GSE157996, generated by our group^8^, along with datasets GSE146981^64^, GSE128033^61^, GSE135893^63^, GSE136831^62^, GSE122960^99^ and GSE132771^100^ from other groups, were all used to determine NAMPT expression in healthy and IPF samples.

The dataset of single cell RNA-seq of mouse epithelial and immune cells are under GSE295739. ScRNA-seq was performed in Cedars-Sinai Medical Center Genomic core. Mouse lung tissue dissociation and single cell isolation were described previously^6^. Mouse AT2s were gated as Epcam^+^CD31^-^CD34^-^CD45^-^CD24^-^Sca-1^-^ population for flow sorting. Sequencing library construction was done by the 10x Genomics chromium platform as previously described^105^. Cell Ranger version 7.0.1 (10x Genomics) was used to process raw sequencing data and Seurat suite version 4.1.0 for downstream analysis. Detailed methods including quality control, cell clustering, doublet calling, and annotation were done as previously described^102^.Differentially expressed genes with a logFC over 0.1 between NAMPT positive (expression > 1) and NAMPT negative (expression ≤ 1) AT2s were analyzed using R software.

#### Bulk RNA-seq and data analysis

Bulk RNA-seq and data analysis were performed as described previously^102^. Detailed methods can be found in Supplementary Methods.

#### Single-cell western blotting

Single-cell western blot analysis for NAMPT was performed following the manufacturer’s instructions (Milo Single Cell Western Blot System). Briefly, single-cell suspensions of healthy and IPF lung AT2s were loaded onto single-cell chips (Standard scWest Kit, K600-1, Proteinsimple) and processed using the Milo system. After buffer washes, the chips were incubated with anti-NAMPT antibody (Cell Signaling Technology, Catalog #61122, RRID: AB_2799602, 1:40 dilution) and anti–β-tubulin antibody (GenScript, clone 2G7D4, Catalog # A01717, RRID: AB_2622210, 1:40 dilution) for 2 hours, followed by incubation with secondary antibodies for 1 hour at room temperature. The chips were then read, and data were analyzed using the Scout system. NAMPT expression was normalized to β-tubulin levels.

### QUANTIFICATION AND STATISTICAL ANALYSIS

The statistical difference between groups in the bioinformatics analysis was calculated using the Wilcoxon Signed-rank test. For the scRNA-seq data the lowest p-value calculated in Seurat was p < 2.2e-10^-16^. For all other data the statistical difference between groups was calculated using GraphPad and the exact value was shown.

## Supplementary Methods

### Human and mouse lung tissue dissociation and flow cytometry

Human lung single cell isolation and flow cytometer were performed as previously described^6^. Flow cytometry was performed with Fortessa and BD Symphony S6 Cell Sorter and analyzed with Flow Jo 10.8.1 software (BD Biosciences). Human AT2 cells were sorted as Epcam^+^CD31^-^ CD45^-^HTII-280^+^ cells. Anti-human CD31 (clone WM59, Catalog # 303118, RRID AB_2247932), anti-human CD45 (clone WI30, Catalog # 304016, RRID AB_314404), anti-human EpCAM (clone 9C4, Catalog # 324212, RRID AB_756086) were from BioLegend. Anti-HTII-280 (Catalog # TB-27AHT2-280, RRID: AB_2832931) were from Terrace biotech.

Mouse lung tissue dissociation and single cell isolation were described previously^6^. Mouse AT2s were gated as Epcam^+^CD31^-^CD34^-^CD45^-^CD24^-^Sca-1^-^ population for f. low sorting. FITC anti-mouse CD24 (Clone M1/69, Catalog #101806, RRID: AB_312839), PE/Cyanine7 anti-mouse Ly-6A/E (Sca-1) (Clone D7, Catalog #108114, RRID: AB_493596) and APC/Cyanine7 Streptavidin (Catalog #108114) were from BioLegend. Biotin anti mouse CD31 (PECAM-1) (Clone 390, Catalog # 13-0311-85, RRID: AB_466421), Biotin anti mouse CD45 (Clone 30-F11, Catalog # 13-0451-85, RRID: AB_466447), Biotin anti mouse CD34 (Clone RAM34, Catalog # 13-0341-85, RRID: AB_466426) were from eBioscience.

### Cell lines

Mouse lung fibroblast cell line MLg2908 (Catalog CCL-206) was from ATCC.

### Bleomycin instillation and bronchoalveolar lavage

Bleomycin instillation was described previously^6^. Under anesthesia, the trachea was surgically exposed. 0.75 – 2.5 U/kg bleomycin (Hospira) in 25 μl PBS was instilled into the mouse trachea with a 25-G needle inserted between the cartilaginous rings of the trachea. Control animals received saline alone. The tracheostomy site was sutured, and the animals were allowed to recover. Mice were sacrificed at different time points and lung tissue were collected for experiments. Bronchoalveolar lavage was described previously^106^, mice were anesthetized, and lungs and heart were surgically exposed. The trachea was cannulated, and the lungs were lavaged three times with 0.8-ml aliquots of cold PBS. The first 0.8-ml bronchoalveolar lavage (BAL) was used for cytokine protein measurement. The live cells from all three 0.8-ml aliquots of BAL were recovered and counted using a hemocytometer. Cytospin preparations of BAL cells were stained conventionally, and differential cell counts were performed.

For mice NAT treatment, the compound NAT (MedChemExpress, Catalog HY-144776) was dissolved in DMSO, mixed with four volumes of Kolliphor (Sigma-Aldrich, Catalog C5135), and further diluted with eight volumes of 5% dextrose (Sigma-Aldrich, Catalog D9434) to prepare the working solution^107^. Mice received daily intraperitoneal injections of NAT or vehicle at a dose of 20 mg/kg for one week, either starting three days before and continuing three days after bleomycin administration or beginning on Day 10 and continuing for seven consecutive doses following bleomycin treatment.

### Hydroxyproline

Collagen contents in mouse lungs were measured with a conventional hydroxyproline method^108^. In brief, lung tissues were vacuum dried and hydrolyzed with 6N hydrochloride acid at 120°C for overnight. Hydroxyproline content was measured and expressed as mg per lung. The ability of the assay to completely hydrolyze and recover hydroxyproline from collagen was confirmed using samples containing known amounts of purified collagen.

### RNA analysis

RNA was extracted using TRIzol reagent. For real-time PCR analysis, 1 μg total RNA was used for reverse transcription with PrimeScript™ RT Master Mix (Perfect Real Time) (TAKARA). 1.25 μl cDNA was subjected to real-time PCR by using Power SYBR Green PCR Master Mix (Applied Biosystems) and the ABI 7500 fast Real-Time PCR system (Applied Biosystems). The specific primers are listing below: Human *NAMPT* forward GCAGAAGCCGAGTTCAACATC, and reverse TTTTCACGGCATTCAAAGTAGGA.

### Western blotting

Proteins were assessed with western blotting as previously described^102^. The membrane were probed with antibodies against human NAMPT (Cell Signaling Technology, Catalog #61122, RRID: AB_2799602, 1:1000), mouse NAMPT (Proteintech, Clone 3D4D8, Catalog #66385-1g, RRID: AB_2881761, 1:1000), human SIRT7 (Santa Cruz, Catolog #sc-365344, RRID: AB_10850175, 1:1000), human SOD2/MnSOD (acetyl K122) (Abcam, Clone NCI-R156-33, Catalog, ab240391, RRID:AB_2892634, 1:1000), human SOD2 (Cell Signaling Technology, Catalog #13194s, RRID: AB_2750869, 1:1000), human HSP60 (ThermoFisher, Catalog #MA3-012, RRID: AB_2121466, 1:1000), human COX IV (Abcam, Catalog #ab202554, RRID: AB_2861351, 1:1000), human Lamin A/C (Cell Signaling Technology, Catalog #2032, RRID: AB_2136278, 1:1000). GAPDH (Cell Signaling Technology, Clone 14C10, Catalog #2118s, RRID: AB_561053, 1:5000) and β-tubulin (GenScript, Clone 2G7D4, Catalog #A01717, RRID: AB_2622210, 1:5000) were used as a loading control.

### Histology and immunofluorescence staining

Freshly dissected human lung tissues were fixed in 10% neutral formalin (Sigma-Aldrich) overnight, dehydrated through an ethanol gradient, and embedded using a tissue embedding machine. Paraffin sections (5 μm thick) were cut using a Microm HM325 microtome, mounted on Superfrost Plus microscope slides, and rehydrated through xylene and an ethanol gradient. Antigen retrieval was performed using citrate-based antigen unmasking solutions before antibody staining. The sections were incubated with anti-NAMPT (Novus, Catalog # NBP2-19467) and anti-HTII-280 (Terrace Biotech, Catalog # TB-27AHT2-280, RRID: AB_2832931) at 1:100 dilution for overnight at 4°C followed by incubation with secondary antibodies (AF488 goat anti-mouse IgM (Jackson Immunoresearch, Catalog #115-545-020, RRID: AB_2338843), Cy3 goat anti-rabbit IgG (Jackson Immunoresearch, Catalog #111-165-003, RRID: AB_2338000)) at 1:400 dilution in blocking buffer for 2h at RT. At least 10 random fields of view per section were photographed (20× objective) on a Zeiss LSM 780 Confocal Microscope (ZEISS). NAMPT signal quantification in HTII-280–positive cells were performed using ZEISS software. Three samples per group were analyzed, with 50–60 individual AT2s measured per lung. For immunofluorescence staining of cells in culture, cells were fixed in 4% paraformaldehyde for 10 min, followed by permeabilization with 0.1% (v/v) Triton-100/PBS for 20 min at RT and blocked with blocking buffer (10%(v/v) goat serum in PBS) at RT for 1h. They were then incubated with primary antibodies, anti-SIRT7 (Santa Cruz, Catalog # sc-365344, RRID: AB_10850175) and anti-TOM20 (Proteintech, Catalog # 11802-1-AP, RRID: AB_2207530), at 1:100 dilution for overnight at 4°C. Subsequently, samples were incubated with secondary antibodies (AF 488 goat anti-mouse IgG (Jackson Immunoresearch, Catalog #115-545-146, RRID: AB_2307324), Cy3 goat anti-rabbit IgG (Jackson Immunoresearch, Catalog #111-165-003, RRID: AB_2338000)) at 1:400 dilution in blocking buffer for 2h at RT. Nuclei were counter-stained with 1 μg/mL DAPI (BD Biosciences, Catalog # 564907, RRID: AB_2869624) for 10 min at RT.

### 3D organoid cultures of human and mouse AT2s

Flow-sorted human (EpCAM^+^HTII-280^+^) or mouse (EpCAM^+^CD24^−^Sca-1^−^) AT2s were cultured in a 1:1 Matrigel/medium mixture in the presence of MLg2908 lung fibroblasts. A total of 3 × 10^3^ AT2s and 2 × 10^5^ MLg2908 cells (Catalog #CCL-206, ATCC) were plated in 100 μl of the Matrigel/medium mix onto 24-well, 0.4 μm Transwell inserts, with 410 μl of medium added to the lower chamber. Cells were cultured either in medium alone or with specified treatments. The medium composition was described previously^6^. Growth factor–reduced Matrigel (Catalog #354230) was obtained from Corning Life Sciences. The following treatments were used: 5 or 10 μM NAT (MedChemExpress, Catalog HY-144776), 5 or 10 μM NAT-5r (Wuxi AppTec), and 5 or 10 μM SBI-797812 (MedChemExpress, Catalog HY-126255); 0.5, and 2 nM FK866 (Sigma-Aldrich, Catalog 481908); 100 μM nicotinamide mononucleotide (NMN, Sigma-Aldrich, Catalog N3501); and 100 μM nicotinamide riboside (NR, Cayman Chemical, Catalog 23132,). 10 mM nicotinamide (NAM, Sigma-Aldrich, Catalog N0636). The same volume of DMSO served as a control. Fresh medium was changed every other day, and cultures were maintained in a humidified incubator at 37°C with 5% CO₂. Colonies were visualized using an Invitrogen™ EVOS™ XL Core Configured Cell Imager (Invitrogen). Colony-forming efficiency (CFE) was determined by counting colonies with a diameter of ≥50 μm per insert and calculating the percentage of input epithelial cells that formed colonies 12 days after plating.

### Bulk RNA-seq and data analysis

Following 48-hour treatment of isolated IPF AT2s with NAT, RNA was extracted as previously described. For NAMPT knockout and activation in immortalized AT2 cell lines, RNA was isolated from cells cultured in 60 mm plates. The extracted total RNA was submitted to the Cedars-Sinai Medical Center Genomics Core and the UCLA Technology Center for Genomics & Bioinformatics (TCGB) for library preparation and RNA sequencing. Sequencing was performed on a NovaSeq X Plus platform, generating 30 million reads per sample. Gene Set Enrichment Analysis (GSEA) of differentially expressed genes was conducted using the GSEA tool^109^. The raw RNA-seq data generated in this study have been deposited in the NCBI Gene Expression Omnibus (GEO) database. (GEO GSE295740).

### Cell growth rate

Cell growth rate of NAT treatment was measured by IncuCyte ZOOM Live Cell Analysis System (Essen BioScience) by following the instruction from the manufacturer.

## QUANTIFICATION AND STATISTICAL ANALYSIS

The statistical difference between groups in the bioinformatics analysis was calculated using the Wilcoxon Signed-rank test. For the scRNA-seq data the lowest p-value calculated in Seurat was p < 2.2e-10-16. For all other data the statistical difference between groups was calculated using Prism (version 10.1.1) (GraphPad Software) and the exact value was shown. Data are expressed as the mean ± SEM. The sample size for in vivo bleomycin fibrosis studies was based on the previous studies in the lab^6^. No animals were excluded for analysis. All experiments were repeated two or more times. Data were normally distributed and the variance between groups was not significantly different. Differences in measured variables between experimental and control group were assessed by using Student’s t-tests. One-way ANOVA followed by Bonferroni’s or Dunnett’s multiple comparison test or two-way ANOVA followed Tukey’s multiple comparison test was used for multiple comparisons. Results were considered statistically significant at p < 0.05.

## Supplemental information

### Supplemental figure legends

**Figure S1.**
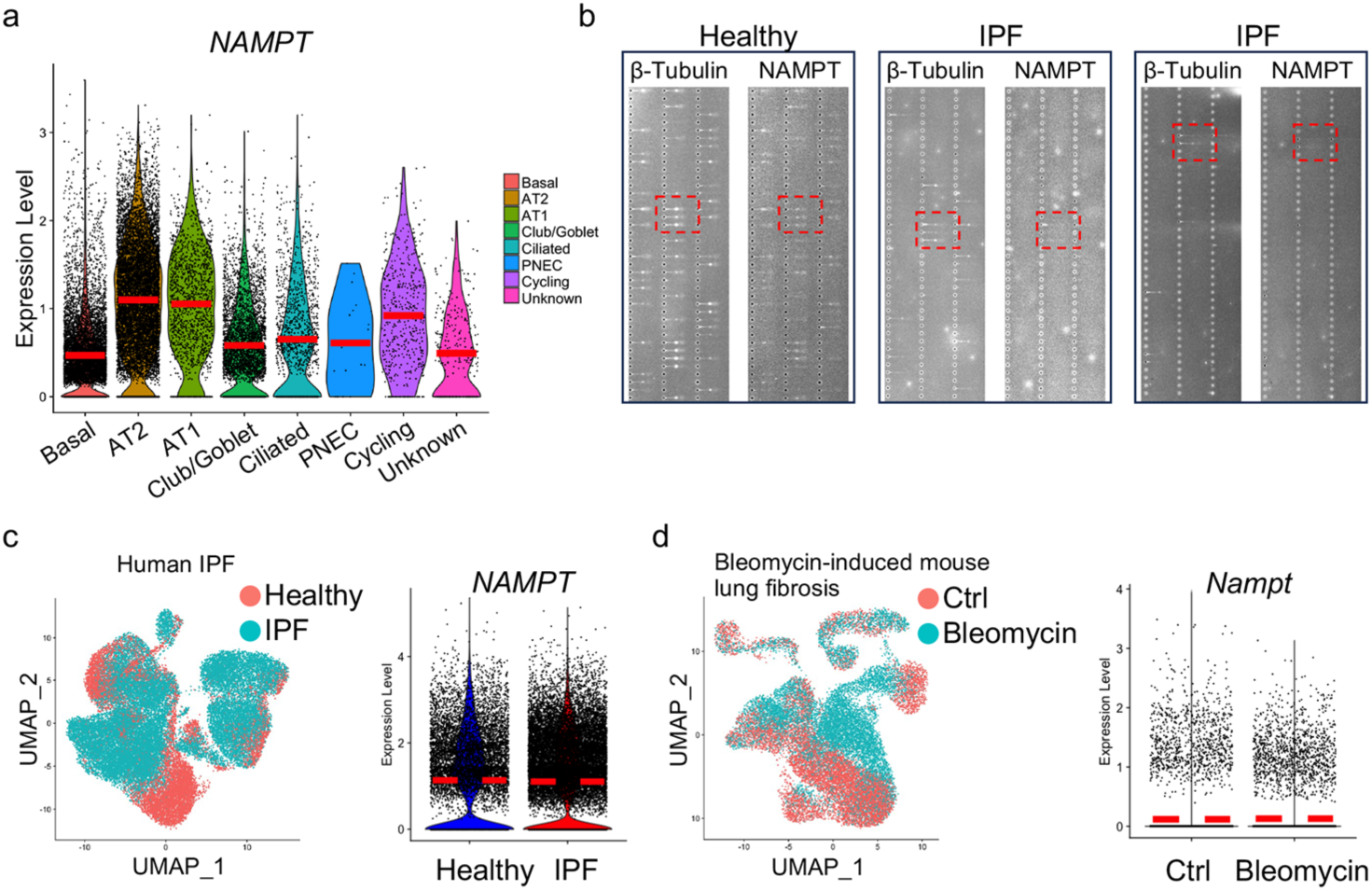
NAMPT expression in epithelial cells and fibroblasts, related to Figure 1. (A) Violin plots showing *NAMPT* expression levels in lung epithelial cell types from in house scRNA-seq dataset. (B) Fluorescence micrographs of single-cell western blot for NAMPT in freshly isolated AT2s from healthy and IPF lungs, with β-Tubulin as a loading control. (C) UMAP visualization of fibroblast populations and *NAMPT* expression in integrated dataset from 7 scRNA-seq datasets on healthy and IPF human lungs (GSE136831, GSE157376, GSE128169, GSE135893, GSE128033, GSE122960, and GSE132771). (D) UMAP visualization of fibroblast populations and *Nampt* expression in integrated dataset of 5 scRNA-seq datasets on bleomycin-induced fibrotic mouse lungs and controls (GSE111664, GSE129605, GSE131800, GSE104154, and GSE132771).

**Figure S2.**
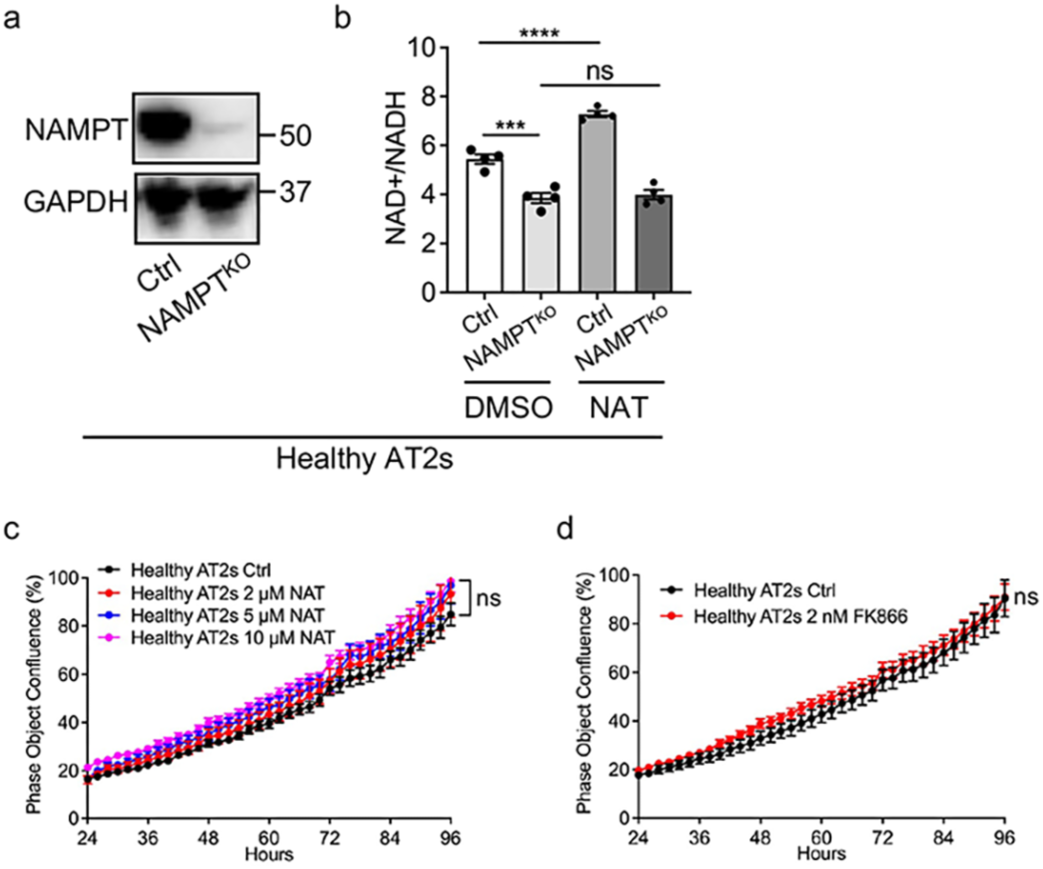
NAT promote NAD^+^ biosynthesis in AT2s specifically through NAMPT, related to Figure 2. (A) Western blot analysis of NAMPT expression in NAMPT^KO^ and control immortalized AT2s. GAPDH served as loading control. (B) NAD^+^/NADH ration in NAMPT^KO^ and control immortalized AT2s with or without NAT treatment (n = 4 per group, ns, no significant, ***p < 0.001, ****p < 0.0001 by ANOVA). (C and D) AT2 cell growth rate after treatment with NAT or FK866 at indicated concentrations (n = 6 per group, ns, no significant by unpaired t tests or ANOVA). Data are shown as the mean ± SEM.

**Figure S3.**
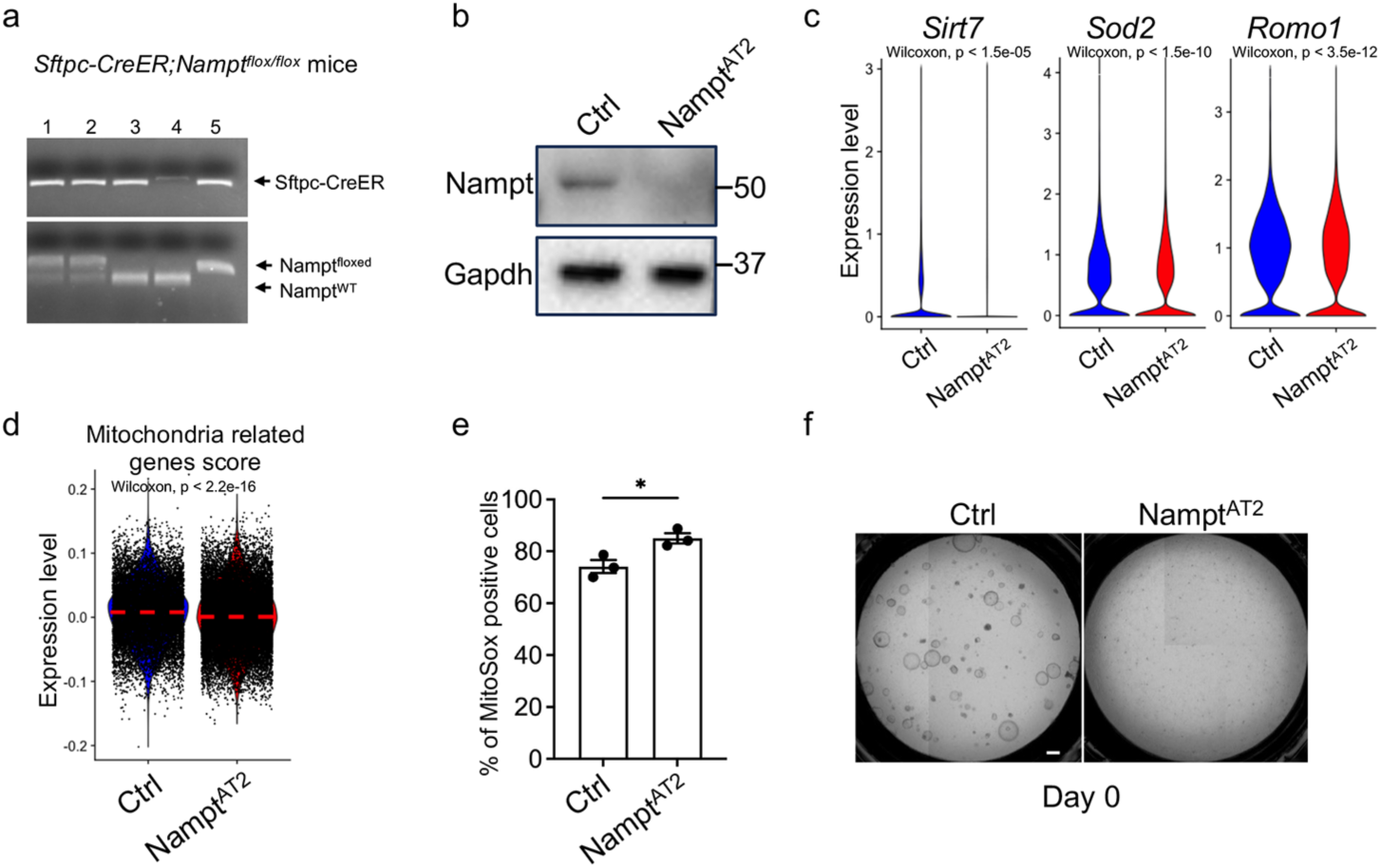
AT2 cell specific Nampt deletion impaired AT2 progenitor renewal in vivo, related to Figure 6. (A) Representative result showed successful genotyping of *Sftpc-CreER*; *Nampf^lox/flox^* mouse in Lane 5. (B) Western blot analysis of Nampt expression in Nampt^AT2^ and control AT2s. Gapdh served as loading control. (C) Violin plots showing expression level of oxidative stress response genes (*SOD2*, *ROMO1*) and *SIRT7*. (D) Mitochondrial related gene score in AT2s from control and Nampt^AT2^ lungs on day 4 after bleomycin injury based on scRNA-seq datasets. (E) Flow cytometry analysis of mitochondrial superoxide levels in AT2s isolated from Nampt^AT2^ and control mice 5 days after bleomycin injury. Data are shown as the mean ± SEM from 3 replicates. *p < 0.05 by unpaired t test. (F) Representative images of colony formation by AT2s from Nampt^AT2^ and control mice without bleomycin injury (D0). Scale bars, 300 μm.

**Figure S4.**
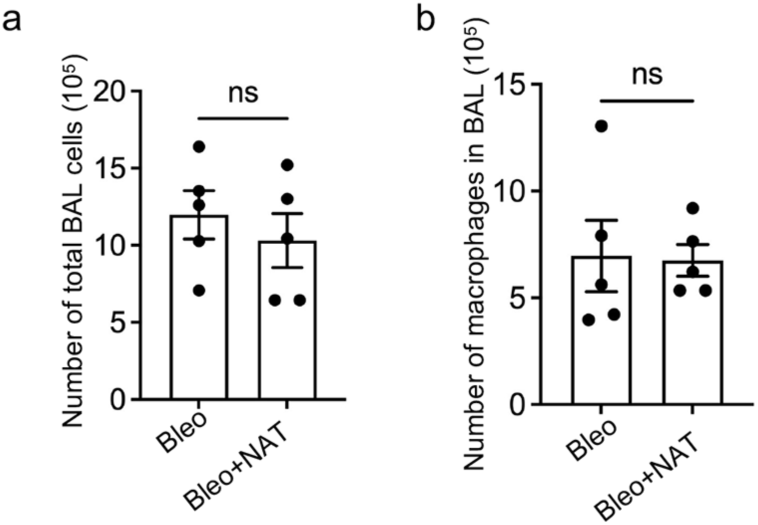
Number of total BAL cells and macrophages in NAT treated and control mice, related to Figure 7. (A) Numbers of total BAL cells and (B) numbers of macrophages in BAL per lung of mice treated with NAT and bleomycin or with bleomycin alone for 5 days as indicated in figure 7A (n = 5 per group). ns, no significant, by unpaired t test. Data are shown as the mean ± SEM.

